# Local inhibitory topology dictates the spatial compartmentalization of hippocampal sharp-wave ripples

**DOI:** 10.64898/2026.06.30.735500

**Authors:** Alexandra Tzilivaki, Daniel Parthier, Annu Kala, Roberto De Filippo, Dietmar Schmitz

**Affiliations:** Charité-Universitätsmedizin Berlin, corporate member of Freie Universität Berlin, Humboldt-Universität Berlin, and Berlin Institute of Health, Neuroscience Research Center, 10117 Berlin, Germany; NeuroCure Cluster of Excellence, Chariteplatz 1, 10117 Berlin, Germany; German Center for Neurodegenerative Diseases (DZNE), 10117 Berlin, Germany; Department of Biology, Humboldt Universität zu Berlin, Berlin 10117, Germany

## Abstract

Hippocampal sharp-wave ripples (SWRs) are essential for memory consolidation and represent among the most synchronous oscillatory events in the brain. Yet, despite their capacity for widespread synchronization, SWRs frequently remain confined to discrete hippocampal domains, revealing a paradox between global coordination and local autonomy. Here, by analysing existing *in vivo* recordings with an experimentally constrained three-dimensional biophysical model, we show that inhibitory activity and inhibitory topology serve fundamentally distinct functions. Whereas perisomatic inhibition gates SWR generation and dendritic inhibition regulates the strength and spectral properties of ripple oscillations, the spatial organization of inhibitory connectivity establishes local computational domains that enable autonomous ripple generators to coexist. Together, our findings identify a spatial dimension of inhibition, in which inhibitory activity governs the emergence and dynamics of SWRs, while inhibitory topology determines their spatial organization and autonomy.

## Introduction

The ability to transform experience into lasting memories relies on the coordinated activity of neuronal populations, with SWRs serving as a principal network signature of memory consolidation^1–5^. Decades of research have illuminated the “clockwork” of SWRs, establishing that coordinated inhibitory activity, particularly from parvalbumin-positive (PV+) interneurons^6–15^, drives the generation and temporal dynamics of these events^9,12,13,15,16^. Yet, this intense focus on *how* SWRs are generated has left a far more fundamental question unaddressed: *where* and *why* do they remain spatially restricted^17^?

Despite SWRs being among the most synchronous events in the mammalian brain^18–21^, analysis on published recordings across the longitudinal axis of the hippocampus reveal that ripple events can remain spatially restricted^17,22,23^ (**Figure 2**). As indicated by previous studies and the presented data (**Figure 2**), SWRs can behave as spatially compartmentalized multifocal events^17^. Qualitatively similar ripples can emerge independently at distinct local loci within the same CA1 network (**Figures 1-2**), even as a subset synchronizes across considerable distances. However, it remains unclear how independent local events coexist alongside spatially widespread synchrony. This paradox suggests that the mechanisms governing *where* SWRs occur are fundamentally distinct from those governing *how* they are generated (**Figure 1**).

**Figure 1:**
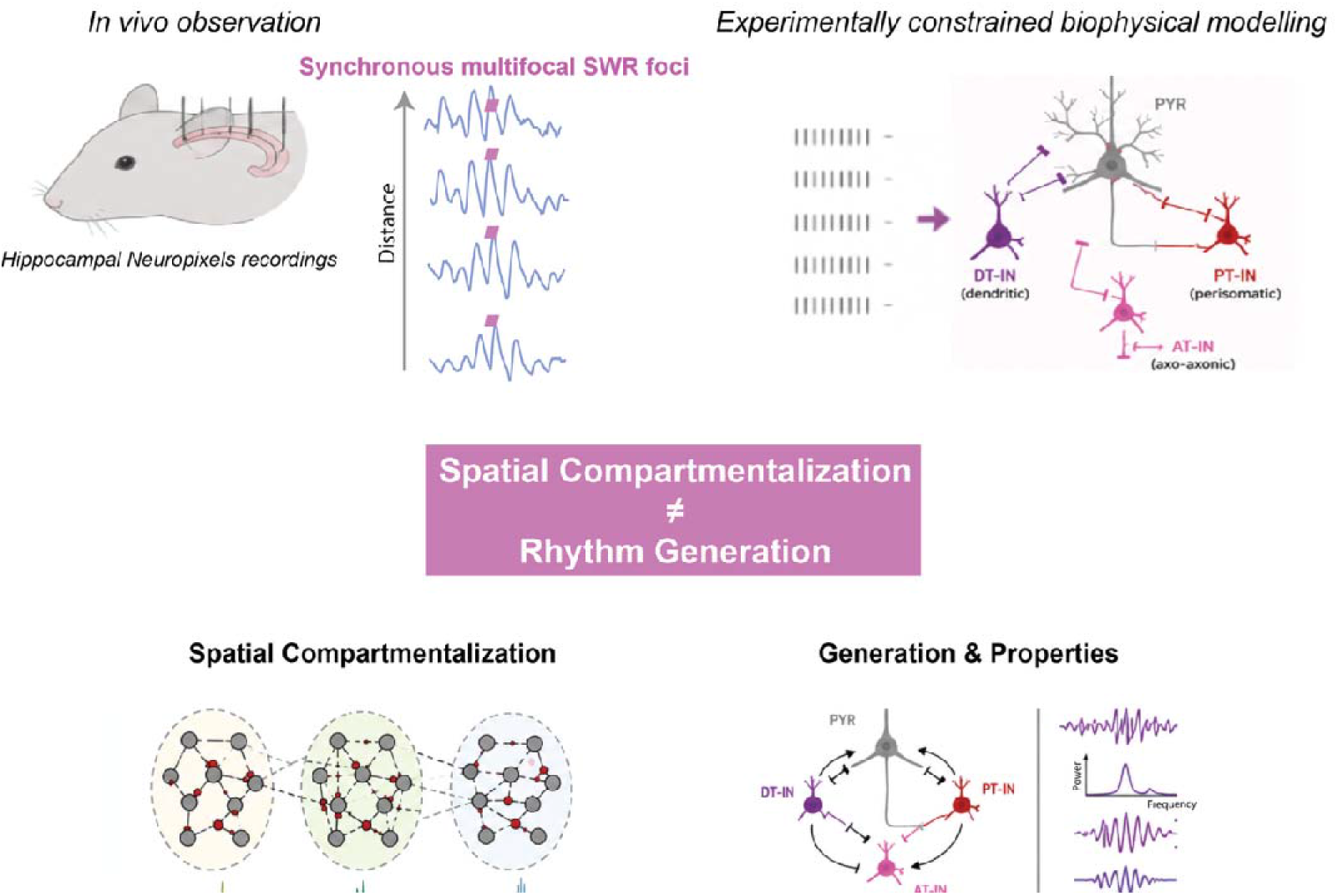
A proposed framework for the spatial organization of hippocampal sharp-wave ripples: inhibitory activity drives generation, inhibitory topology dictates spatial compartmentalization. The central paradox of hippocampal CA1 SWRs: in vivo high-density Neuropixels recordings (upper left) reveal that highly synchronous SWRs can also remain spatially restricted as multifocal events, rather than always propagating globally (data adopted from ^28^. The experimentally constrained, three-dimensional biophysical model of the CA1 network (upper right). The model explicitly incorporates local spatial geometry, distance-dependent synaptic connectivity, and three distinct populations of interneurons: perisomatic-targeting (PT-IN), dendritic-targeting (DT-IN), and axo-axonic (AT-IN) cells. The proposed conceptual framework (middle and bottom panels): Inhibitory topology partitions the network into autonomous functional domains (spatial compartmentalization), whereas inhibitory activity acts as the temporal engine driving the oscillation (rhythm generation).

**Figure 2:**
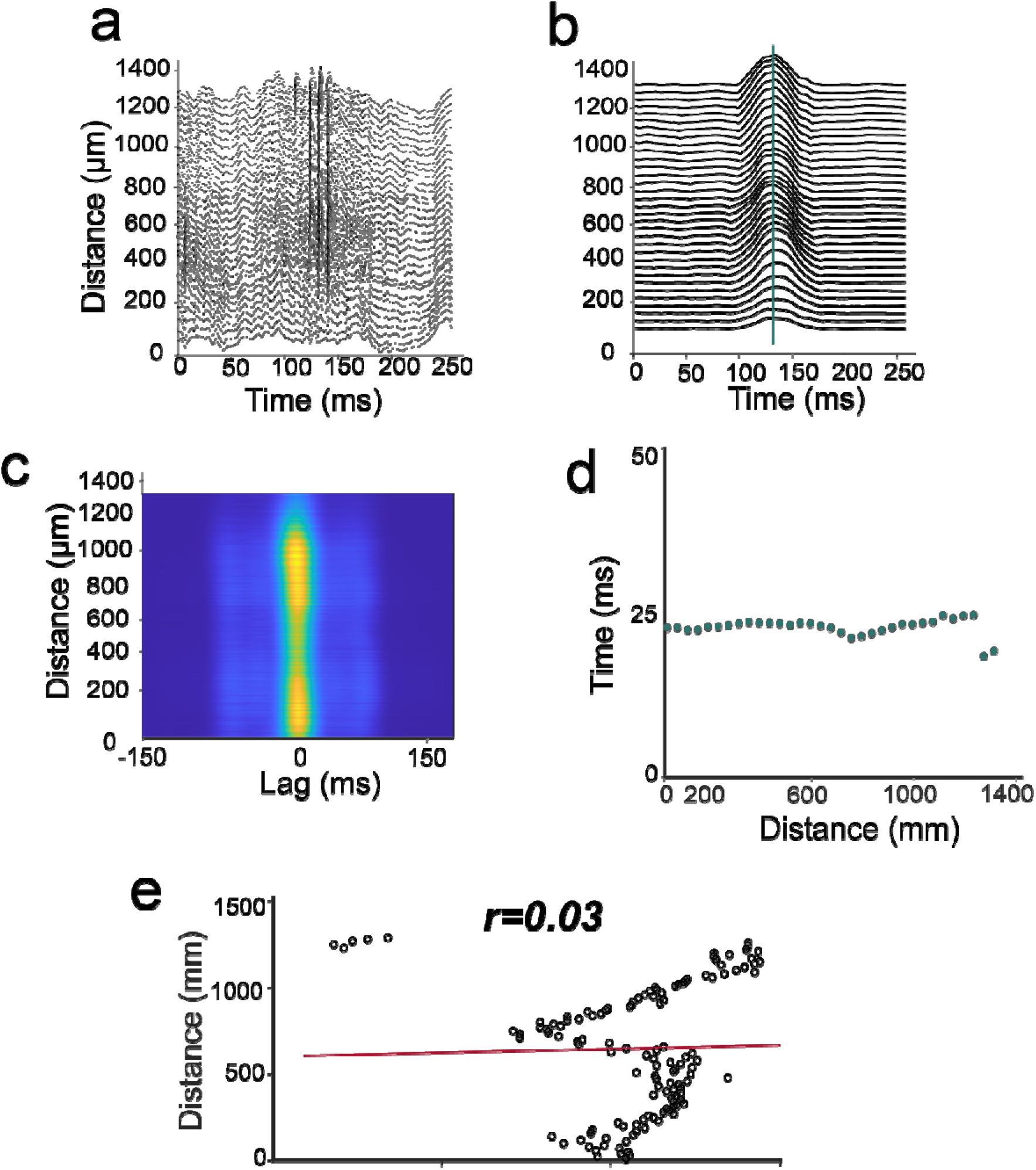
Neuropixels datasets reveal multiple local SWR foci. **(a–b)** Published In vivo Neuropixels recordings along the longitudinal axis of the CA1 region in awake, head-fixed mice, adopted and reanalysed from ^28^. Representative simultaneous LFP recordings across 33 recording sites spanning ∼1.3 mm of hippocampal tissue reveal SWR events that remain tightly spatially confined despite strong temporal synchrony. Representative traces of the raw LFP (a) and envelope of the Hilbert-transformed ripple-band filtered signal (b). **(c)** Cross-correlation heatmap of the Hilbert-transformed ripple envelope between channel 1 and all other channels. **(d)** Time points of the peak Hilbert-transformed ripple power plotted as a function of spatial position, revealing a highly localized spatial footprint without a systematic temporal shift. **(e)** Linear regression analysis between peak ripple timing and electrode position, indicating a lack of consistent temporal progression across the recording axis (Pearson’s r = 0.03, p < 0.0001, Student’s t-test). The stability of ripple peak timing across distance indicates that these events represent stationary, autonomous local generators rather than directional traveling waves. Plots were created from the dataset in ^28^ .

We hypothesize (**Figure 1**) that this spatial compartmentalization is dictated not by inhibitory activity alone, but by the topology of inhibitory connectivity. In biological systems, “small-world” network architectures resolve similar paradoxes by preserving strong local clustering while maintaining network-wide flexibility^24–27^. In the hippocampus, axonal projections are spatially constrained. According to Peters’ rule, synaptic connectivity probability decreases with distance, meaning interneurons are substantially more likely to interact with neighboring cells than distant targets^26^. We challenge the implicit assumption that interneuron activity is merely a temporal pacemaker. Instead, we propose a dual function: the same local interneuron microcircuits that generate the rhythm simultaneously function as a structural blueprint, physically partitioning the CA1 network into partially independent functional domains.

To test this hypothesis, we built an experimentally constrained, three-dimensional biophysical model of the CA1 network (**Figures 3-8**). Crucially, by explicitly incorporating realistic spatial geometry and distance-dependent connectivity rules, our model overcomes the limitations of previous frameworks and allows us to directly disentangle inhibitory activity from its spatial architecture (**Figure 3**). By systematically manipulating network topology, we predict that while interneuron activity acts as the rhythm generator (**Figures 4,6,7**), structured inhibitory topology is the essential architect of SWR spatial compartmentalization, enabling multiple autonomous computational units to coexist (**Figures 5,8**).

**Figure 3.**
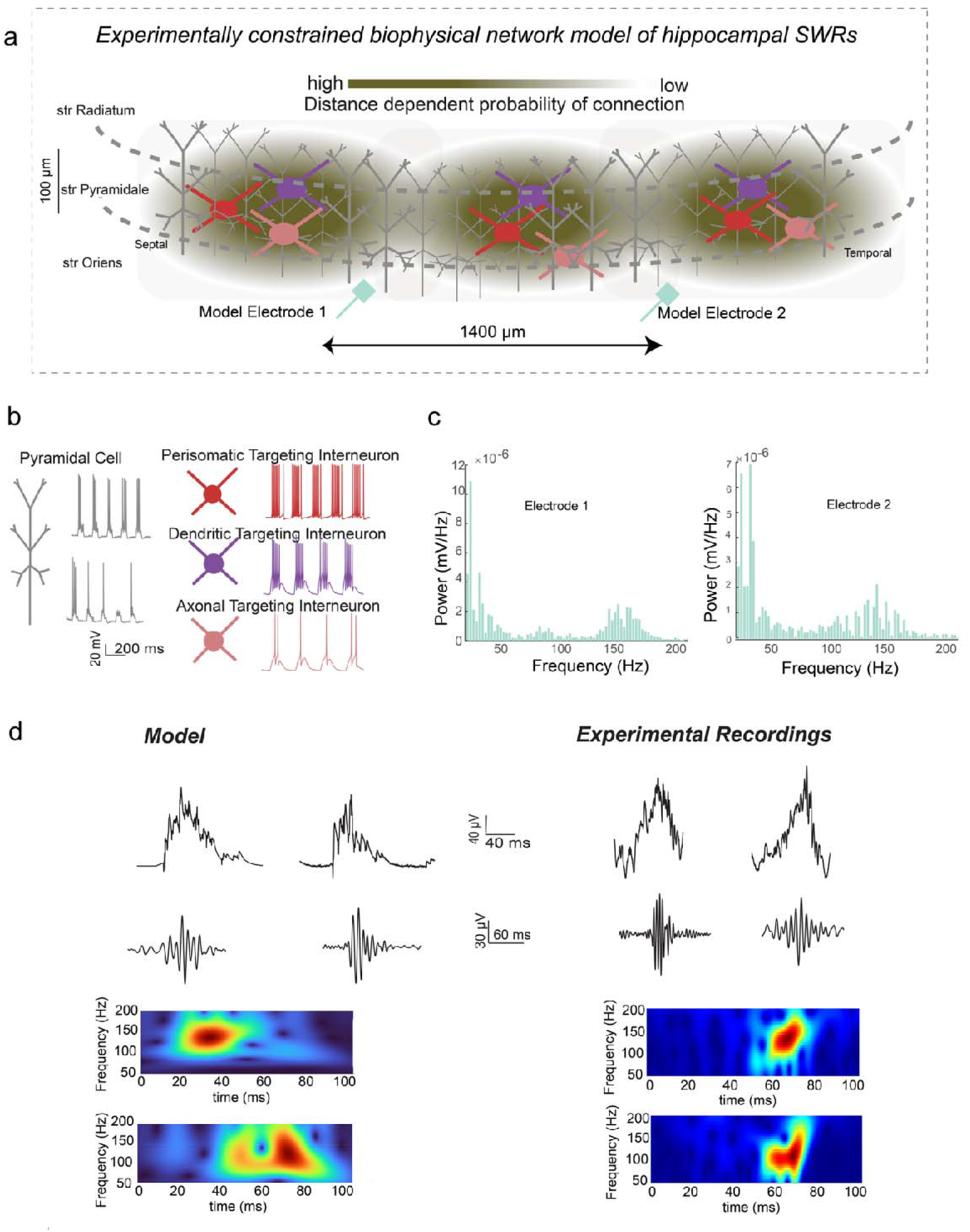
An experimentally constrained topological biophysical model of CA1 reproduces local SWRs. **a)**Schematic of the three-dimensional, multicompartmental CA1 network architecture. Pyramidal cells and three distinct inhibitory interneuron populations are distributed across their anatomically correct laminae (stratum oriens, pyramidale, and radiatum). Synaptic connectivity follows Peters’ rule and experimental data, utilizing a distance-dependent decay probability (green gradient) to naturally establish a structured local topology. Two virtual extracellular electrodes (Model Electrode 1 and 2) are spaced 1400μm apart to mirror our in vivo Neuropixels configuration. **(b)** Morphological representations and simulated physiological firing patterns of the principal pyramidal cells and the three modeled inhibitory interneuron subpopulations: perisomatic-(red), dendritic-(purple), and axonal-targeting (pink) cells. The firing activity of each population during simulated SWRs is robustly constrained by experimental evidence: pyramidal cells typically fire 1–3 spikes per event, while perisomatic, dendritic, and axonal interneurons fire 7–8, 4–6, and 1–2 spikes per event, respectively. **(c)** Welch power spectral density (PSD) estimates of the simulated LFP recorded at both virtual electrodes. The spectra reveal the classical, bimodal electrophysiological phenotype of SWR: high power in the low-frequency (1-50 Hz) sharp-wave band coupled with a distinct, prominent peak in the high-frequency ripple band (∼100–200 Hz). **(d)** Direct electrophysiological comparison of in silico simulated SWRs (left) against our in vivo experimental recordings (right). Top traces display the raw LFP, capturing the pronounced low-frequency sharp-wave deflection. Middle traces show the identically processed, band-pass filtered signals isolating the high-frequency ripple oscillation (110–250 Hz). Bottom panels present time-frequency spectrograms of the corresponding events. The model faithfully captures the transient temporal envelope, the precise spectral power distribution, and the characteristic physiological trial-to-trial variability of biological SWRs. (Note: The simulated neuronal firing traces, spectra, and LFPs shown in panels **b**, **c**, and **d** serve as representative examples drawn from many independent, simulation trials).

**Figure 4.**
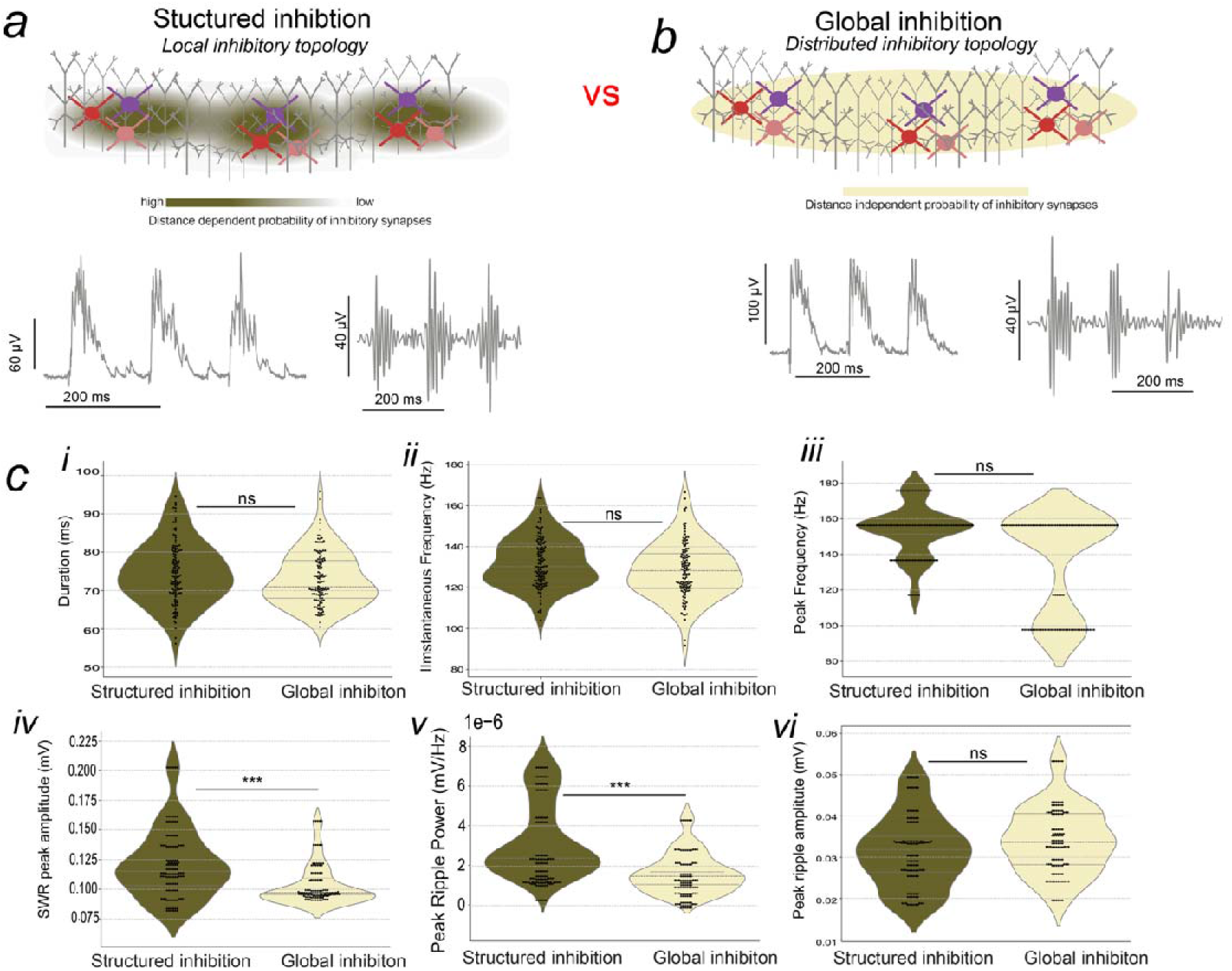
Structured and global inhibitory topologies both support SWR generation. **(a, b)** Network schematics and representative simulated recordings contrasting the biologically realistic structured topology against an engineered global inhibitory topology. **(a)** Under structured inhibition, inhibitory synaptic connectivity follows a distance-dependent probability rule (green gradient), anchoring interneurons to their local microcircuits. **(b)** Under global inhibition, interneuron outputs are spatially randomized (distance-independent uniform distribution, yellow), abolishing local topological boundaries while preserving the underlying distance-dependent excitatory architecture. Representative raw LFP traces (left, sharp waves) and band-pass filtered traces (right, ripples) demonstrate that homogeneous CA3 drive elicits robust, physiologically realistic SWRs in both network configurations. **(c)** Comprehensive quantification of SWRs temporal and spectral properties across the two network topologies. Violin plots compare event duration (i), instantaneous ripple frequency (ii), peak ripple frequency (iii), SWR raw peak amplitude (iv), peak ripple power (v), and peak ripple amplitude (vi). The global topology fully retains the fundamental capacity to generate the SWR rhythm, preserving core temporal features including event duration and oscillatory frequency (ns, not significant). However, the loss of local inhibitory clustering triggers a significant reduction in overall event magnitude, explicitly dampening SWR peak amplitude (p < 0.001) and peak ripple power (p < 0.001). Thus, while structured inhibitory topology modulates the maximum strength of the oscillation, it is not a prerequisite for rhythm generation itself. Data represent n = 120 structured SWR events and n = 120 global SWR events, extracted across 10 independent, randomized simulation trials per condition. Statistical significance was assessed using generalized linear mixed-effects model to rigorously account for hierarchical trial variance. Boxplots show mean and std values.

**Figure 5.**
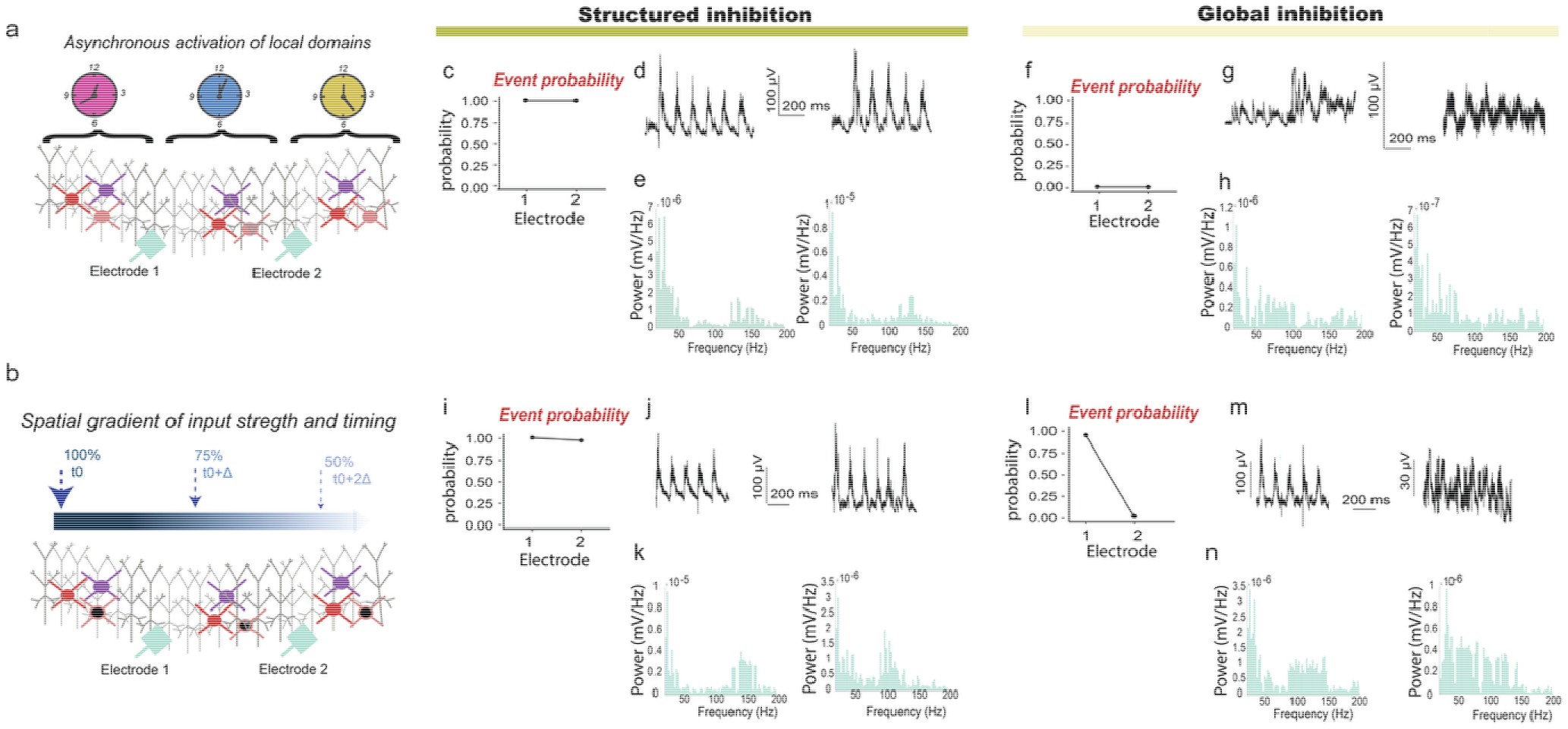
Structured inhibitory topology preserves local SWR foci under heterogeneous input conditions. (**a, b**) Schematics of the in silico spatiotemporal stress tests. (**a**) Protocol A (Asynchronous activation) introduces temporally different inputs to different thirds of the CA1 axis, creating a severe network-wide temporal conflict. (**b**) Protocol B (Spatiotemporal gradient) progressively decreases input strength (100% to 50%) while increasingly delaying onset across the longitudinal axis, creating a stark spatial imbalance. (**c–e, i–k**) Under structured inhibition, the network acts as an “architectural firewall”. (**c**-**e**) Despite severe temporal mismatch (Protocol A), spatially constrained inhibitory motifs isolate neighboring domains, maintaining ∼100% SWR event probability (**c**) and robust electrophysiological SWR signatures (**d, e**) at both recording sites. (**i-k**) Under spatial imbalance (Protocol B), structured inhibition prevents dominant domains from hijacking the network; even the spatial pole receiving 50% weaker, delayed input (Electrode 2) successfully generates autonomous SWRs matching the physiological signature of the strongly driven pole. (**f–h, l–n**) Under global inhibition, the unconstrained topology leads to network collapse. (**f-h**) Confronted with temporal mismatch (Protocol A), the lack of spatial constraints disrupts the ability of the network to maintain coordinated SWR generation. Asynchronous activation cascades widespread inhibition across the entire network, destroying precise synchronization and completely abolishing SWR generation at all sites (event probability = 0). (**l-n**) Confronted with a spatial gradient (Protocol B), global inhibition forces destructive, winner-takes-all competition. While SWRs survive at the pole receiving the strongest, earliest input (Electrode 1), this dominant domain broadcasts globalized inhibition that actively and completely suppresses the domain receiving weaker, delayed excitation (Electrode 2). Thus, we predict that local inhibitory architecture is a critical structural requirement to preserve independent memory-related computations during heterogeneous network states.

**Figure 6.**
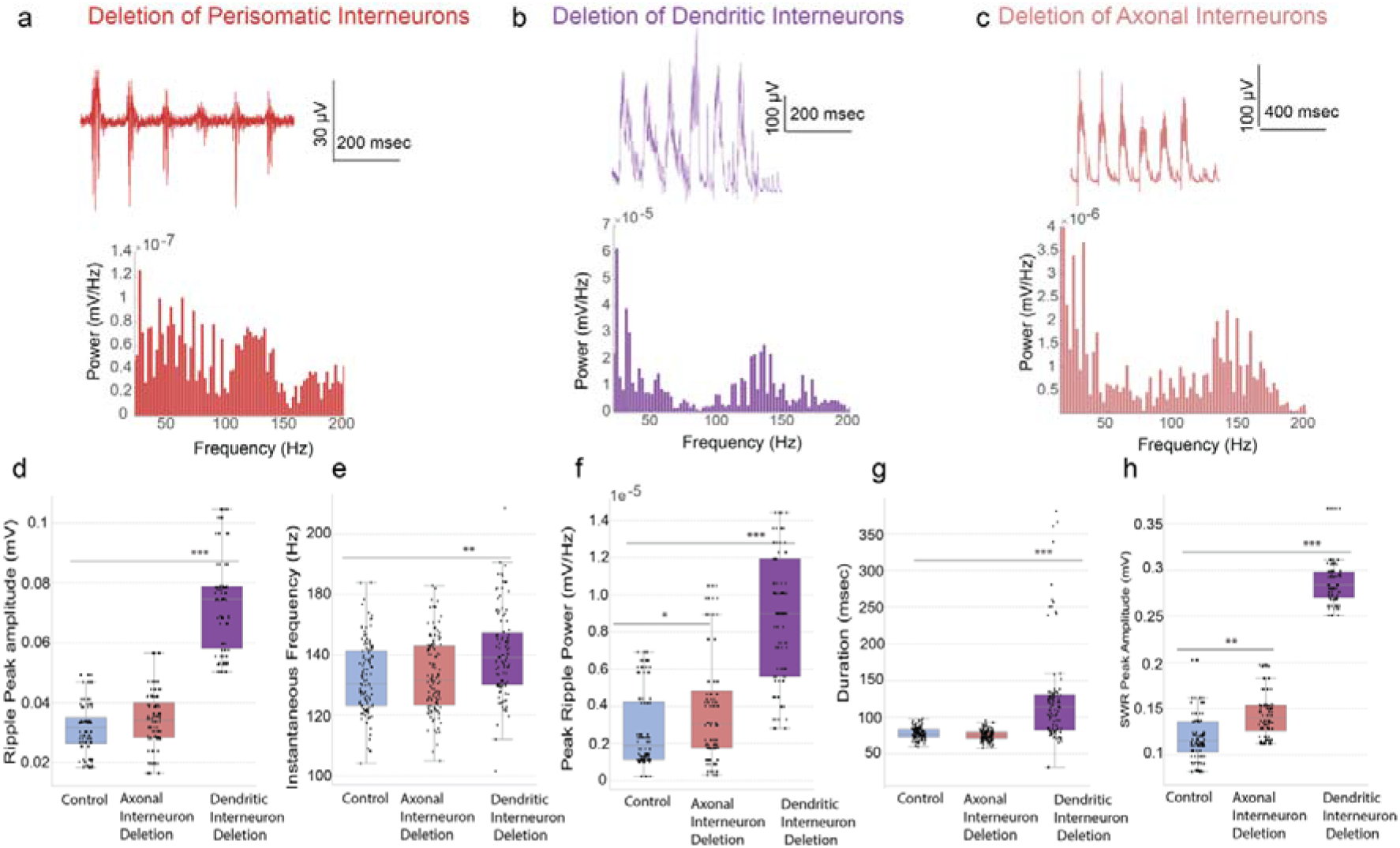
Selective deletion of interneuron populations differentially alters SWR dynamics. **(a–c)** Representative simulated LFP traces (top) and corresponding Welch power spectral densities (bottom) following targeted in silico ablation of specific inhibitory interneuron populations. **(a)** Deletion of perisomatic-targeting interneurons acts as a binary gate, abolishing SWR generation and leaving unstructured, low-amplitude spiking activity. **(b)** Conversely, deletion of dendritic-targeting interneurons triggers runaway pyramidal excitation, resulting in high-amplitude sharp-wave deflections and elevated ripple power. **(c)** Deletion of axonal-targeting interneurons produces only minor spectral alterations, preserving baseline SWR dynamics. (Note: LFP traces and spectra shown are representative examples from independent, randomized simulation trials). **(d–h)** Quantification of SWRs electrophysiological features across the targeted deletion conditions. Box plots and overlaid individual data points compare ripple peak amplitude (**d**), instantaneous ripple frequency (**e**), peak ripple power (**f**), event duration (**g**), and overall SWR raw peak amplitude (**h**). Ablation of dendritic inhibition inflates all measured magnitude and temporal parameters, explicitly defining its computational role as a continuous gain-control mechanism that restricts the participating excitatory ensemble. Data represent n = 120 analyzed SWR events per condition, extracted across multiple independent, randomized simulation trials. Statistical significance was evaluated using generalized linear mixed-effects models to account for hierarchical trial variance (*p < 0.05, **p < 0.01, ***p < 0.001). Boxplots show mean and std values.

## Results

### In vivo Neuropixels data reveal spatially segregated SWR foci

To map the spatial architecture of hippocampal SWRs, we analysed published *in vivo* Neuropixels recordings along the longitudinal axis of the CA1 region from head-fixed mice moving on a treadmill^28^. This configuration enabled simultaneous, high-resolution recording of local field potential (LFP) activity across the hippocampal formation, capturing SWR activity from approximately 130 channels, encompassing ∼1.3 mm of hippocampal tissue (**Figure. 2a–c**).

While SWRs are classically viewed as highly synchronous events capable of engulfing the entire network, our in vivo data analysis revealed a different reality. We observed SWR events that, despite exhibiting strong temporal synchrony, remained tightly spatially confined. Rather than recruiting the entire CA1 axis, these events highlight the existence of isolated, local ripple foci operating within the broader network (**Figure. 2a–c**).

To distinguish whether these confined events were merely propagating waves or truly independent stationary generators, we quantified the spatiotemporal footprint of the ripple activity. We extracted the timing of the peak Hilbert-transformed ripple envelope from each channel and mapped it against the spatial position of its corresponding electrode (**Figure. 2d–e**). If these SWRs were traveling waves, we would expect a systematic temporal delay across the recording sites, reflecting a directional sweep of activity along the CA1 axis.

Instead, the SWRs analyzed here exhibited no consistent temporal progression across the ∼1.3 mm of recorded tissue. The timing of peak ripple power was remarkably stable across the recording sites, and linear regression analysis indicated an almost complete absence of spatial-temporal correlation (Pearson’s r=0.03, **Figure. 2f**). These findings provide *in vivo* evidence that temporally coordinated SWRs can remain firmly anchored in space, supporting the existence of multiple autonomous ripple generators. Importantly, these same dataset also contain SWRs that propagate along the longitudinal axis (as we show in^28^), consistent with earlier reports of ripple propagation^23^. In the present study we focus on the spatially segregated regime and the mechanism that allows independent local generators to coexist. This observation raises a fundamental question: how can highly synchronous oscillatory events remain locally restricted within the same hippocampal network? This paradox motivated our hypothesis that the mechanisms governing the spatial compartmentalization of SWRs are distinct from those underlying their generation.

### An experimentally constrained biophysical network model recapitulates the spatial and electrophysiological features of local sharp-wave ripple activity

Our central hypothesis is that while the generation of SWRs depends on the coordinated activity of neuronal populations, their spatial organization may emerge from the underlying topology. Although previous computational models have provided fundamental insights into the cellular and synaptic mechanisms underlying SWR generation^29,30^, they do not incorporate the three-dimensional architecture of CA1 and the distance-dependent constraints that shape neuronal connectivity.

To overcome these limitations, we developed a multicompartmental biophysical model of the rodent CA1 region incorporating experimentally reported anatomical^14,26,31^, electrophysiological^32–37^, and synaptic properties^32,38–40^ (**Figure 3a,b,c**). Neurons were positioned in a realistic three-dimensional CA1 geometry, preserving its characteristic laminar organization and longitudinal “banana-shaped” structure. Pyramidal cell somata were localized within stratum pyramidale, while interneuron populations were distributed according to their experimentally reported anatomical locations. Importantly, connectivity was not assigned randomly but followed experimentally measured connection probabilities and a distance-dependent rule based on Peters’ principle and experimental evidence demonstrating a higher probability of synaptic interactions between nearby neurons^26,41^ (see **Methods**). Given that typical axonal arborizations of pyramidal neurons and interneurons extend approximately 600–900 μm, neurons are expected to preferentially interact with neighboring cells within their anatomical reach. This generated a structured network topology in which local interactions were favored while preserving the possibility of longer-range connections. Therefore, the model does not impose artificially isolated compartments but rather captures the gradual spatial decay of connectivity observed in biological networks.

To reproduce the cellular diversity known to participate in SWRs^42–47^, the model included pyramidal cells and the major interneuron populations that have been experimentally shown to fire during ripple events^10,38–40,48–51^, namely perisomatic-targeting (aka PV+ fast spiking basket cells), dendritic-targeting (a.k.a bistratified) cells, and axonal targeting (a.k.a chandelier cells) (**Figure 3b**). All neuronal models were validated against their experimentally reported electrophysiological properties and synaptic connectivity profiles (**Methods and Supplementary Tables 1–4**). To directly compare the model with our *in vivo* recordings (**Figure 2**), we performed simulated multi-electrode recordings across the longitudinal axis of the network, with virtual electrodes positioned at distant locations along the CA1 structure. The network received a rhythmic excitatory drive mimicking Schaffer collateral input from CA3 together with stochastic background activity to reproduce the natural fluctuations of physiological states (**Figure 3c, d**). Under these conditions, the model spontaneously generated characteristic SWR events consisting of a low-frequency sharp-wave component (1–50 Hz)^5^ and a high-frequency ripple oscillation (80–250 Hz) as per ^18,52^ recorded across both virtual electrodes (**Figure 3c**). Importantly, the generated events were not stereotypical but exhibited trial-to-trial variability in their spectral and temporal characteristics, including sharp-wave amplitude, ripple amplitude, ripple power, peak and instantaneous ripple frequency, and event duration (**Supplementary Figure 1**), consistent with the variability observed in experimental recordings ^18,52,53^.

Beyond reproducing extracellular signatures, the model accurately captured the firing behavior of pyramidal cells^42,45,46,54^ and individual interneuron populations during SWRs, including the number of spikes per event reported experimentally (**Figure 3b** and **Supplementary Figure 2**). To ensure that these responses were not the consequence of excessive parameter tuning, we performed extensive sensitivity analyses by systematically increasing and decreasing both the external CA3 input strength and intrinsic CA1 synaptic conductances by 20–30%. SWRs generation remained stable across these perturbations (**Supplementary Figure 3**), demonstrating that the model dynamics emerge robustly from biologically realistic network organization rather than from a narrow parameter space. Finally, because experimental pharmacological blockade of GABA receptors using gabazine abolishes SWR generation in hippocampal preparation^11,13^, we removed inhibitory synaptic transmission in the model and observed a complete loss of SWR activity (**Supplementary Figure 4**), further validating the ability of the model to reproduce key experimental observations.

Having established this robust, biologically realistic framework, we next leveraged it to disentangle inhibitory activity from inhibitory topology.

### SWR generation is preserved across structured and global inhibitory topologies

Having established a biologically realistic model that faithfully reproduces hippocampal SWRs, we next used this framework to isolate the contribution of inhibitory connectivity topology. To this end, we generated a second network version in which the distance-dependent organization of inhibitory connections was removed (**Figure 4a-b**). Importantly, all other aspects of the model remained identical to the structured network, including the three-dimensional CA1 architecture, cellular composition, intrinsic electrophysiological properties, synaptic strengths, and the distance-dependent connectivity of excitatory pyramidal neurons.

In the global inhibitory network, inhibitory neurons were no longer restricted by spatial proximity and could establish synaptic contacts with both pyramidal neurons and other interneurons anywhere along the longitudinal axis of CA1 (**Figure 4b**). Thus, while the total number of inhibitory synaptic contacts remained unchanged, the identity of their postsynaptic targets became spatially unconstrained. In contrast to the structured network (**Figures 3a, 4a**), where inhibitory neurons preferentially interact with neighboring cells according to anatomical distance constraints, the global model allows stochastic long-range inhibitory interactions. Importantly, this architecture resembles the assumptions of many previous computational SWR models, where spatial constraints of connectivity are generally not explicitly incorporated.

Following activation with the same CA3-like rhythmic input and stochastic background activity used for the structured model, the global inhibitory network also generated robust SWR events (**Figure 4**). Visual inspection of representative LFP traces revealed highly similar sharp-wave and ripple events between the two network architectures. Quantification of SWR spectral properties across multiple simulation trials and recordings from both virtual electrodes demonstrated that the majority of SWR characteristics were preserved despite the absence of local inhibitory topology (**Figure 4c**). Although structured inhibition produced slightly larger ripple power and SWR amplitude compared with the global inhibitory configuration, parameters including ripple frequency and event duration remained largely unaffected.

Consistent with these findings, the firing activity of pyramidal cells and interneurons was comparable between structured and global inhibitory networks (**Supplementary Figure 5**), indicating that inhibitory topology does not fundamentally alter the recruitment of neuronal populations during SWRs.

Together, these findings demonstrate that the emergence of SWRs does not depend on the spatial organization of inhibitory connectivity. While local inhibitory architecture can modulate certain spectral properties of the oscillation, the generation of the rhythm itself is preserved even when inhibitory interactions become spatially global. These results suggest that the primary role of structured inhibition may not lie in determining whether SWRs occur, but rather where they occur across the hippocampal network.

### Structured inhibition enables the coexistence of multiple local SWR foci

For multiple local SWR foci to coexist, individual CA1 subdomains must be capable of generating ripple activity independently, shielded from fluctuations occurring at distant locations along the longitudinal axis. This functional compartmentalization provides a mechanistic explanation for our *in vivo* observations (**Figure 2**), where spatially separated recording sites exhibit independent SWR events despite belonging to the same hippocampal network.

To directly test whether inhibitory topology dictates this spatial independence, we stress-tested our networks with heterogeneous inputs. While previous simulations utilized identical synchronous activation, the true test of local autonomy is whether an individual region can maintain ripple generation when neighboring domains receive conflicting temporal or synaptic drives.

We first designed a temporally heterogeneous “clock” protocol (**Figure. 5a**), in which different thirds of the CA1 model received inputs of identical strength but progressively delayed onset times. This manipulation introduced strict temporal disagreement between neighboring regions. Remarkably, the structured inhibitory network easily absorbed this conflict, remaining fully capable of generating SWRs at both recording sites (**Figure. 5c–e**). Spatially constrained inhibitory motifs isolated neighboring domains from irrelevant temporal fluctuations, allowing each subnetwork to synchronize to its own incoming drive and maintain an independent ripple generator.

In striking contrast, this exact same temporal mismatch completely abolished SWRs in the global inhibitory network (**Figure. 5f–h**). Because interneurons were untethered from their local environment, temporally mismatched activation in one subdomain cascaded widespread inhibition to regions that were either not yet activated or already active. This created a network-wide temporal conflict that destroyed the precise synchronization required for ripple emergence. Under a global topology, local variability rapidly cascades into global instability.

We next asked whether structured inhibition could preserve local ripple generation under a spatiotemporal “gradient” protocol (**Figure. 5b**), progressively decreasing input strength by up to 50% across the longitudinal axis while simultaneously delaying its onset. Again, the structured network proved remarkably robust. SWRs were preserved not only at the pole receiving the strongest input, but also at the opposite pole receiving weaker and delayed excitation (**Figure. 5i–k**).

Conversely, the global inhibitory network collapsed into competition. While SWRs survived near the region receiving the strongest and earliest input, ripple activity was entirely suppressed in the domain receiving weaker excitation (**Figure. 5l–n**). Globally distributed inhibitory interactions allowed the dominant region of the network to stymie weaker local generators, preventing their coexistence.

Together, these stress tests demonstrate that inhibitory topology does not simply regulate whether SWRs can be generated; it determines whether multiple SWR generators can coexist. Structured inhibition creates spatially autonomous computational domains, whereas global inhibition forces destructive competition. Having established that local topology governs *where* SWRs emerge, we next investigated the specific cellular inhibitory mechanisms that dictate *how* these oscillations are generated and shaped.

### Perisomatic inhibition gates generation, whereas dendritic inhibition shapes SWR dynamics

Although the hippocampus contains a highly diverse repertoire of inhibitory interneurons^9^, their specific contributions to SWR generation and modulation remain incompletely understood. A substantial body of experimental evidence has established a critical role of PV+ interneurons in SWR activity, both in *vitro* and *in vivo*^10,12,13,27,32,55–58^. However, PV+ interneurons do not represent a homogeneous population, as distinct subtypes target different subcellular compartments of pyramidal neurons and exhibit distinct firing patterns during SWRs. Therefore, our biophysical model incorporated three major PV+ inhibitory populations: perisomatic-targeting basket cells, dendritic-targeting bistratified cells, and axo-axonic chandelier cells targeting the axon initial segment. This approach enabled us to overcome current experimental limitations and systematically dissect the contribution of distinct inhibitory domains to hippocampal SWR dynamics.

Selective removal of perisomatic inhibition resulted in a complete loss of SWR generation (**Figure 6a**), further confirming the indispensable role of fast perisomatic inhibition in establishing coordinated ripple oscillations. Importantly, this requirement was preserved in both structured and global inhibitory topology network configurations (data not shown), demonstrating that the necessity of perisomatic inhibition for SWR generation is independent of the spatial organization of inhibitory connectivity.

In contrast, removal of dendritic inhibition did not abolish SWRs but profoundly altered their spectral characteristics. SWR events displayed approximately a threefold increase in raw amplitude (**Figure 6b, Supplementary Figure 7**), accompanied by increased ripple peak amplitude and ripple power (**Figure 6d,f**). Additionally, SWRs exhibited longer durations and higher instantaneous ripple frequencies (**Figure 6g,e**). Silencing dendritic targeting interneurons resulted in similar effects in the global inhibitory topology network (**Supplementary Figure 7**). Therefore, while dendritic inhibition is not essential for the existence of SWRs, it acts as a major regulator of their intensity and temporal dynamics.

Removal of axo-axonic inhibition produced comparatively minor effects (**Figure 6c**). Given the relatively low firing rates of chandelier cells during SWRs^32^, both experimentally and in our model (**Figure 1b; Supplementary Figure 2**), the majority of SWR features remained similar to control conditions, with only a modest increase in ripple power (**Figure 6f**).

To identify the cellular mechanisms underlying these changes, we examined the firing activity of individual neuronal populations during SWRs (**Supplementary Figure 6**). Removal of dendritic inhibition resulted in a pronounced increase in pyramidal cell firing, reaching approximately threefold higher spike rates compared to control conditions. This increase closely paralleled the enhancement in SWR amplitude, suggesting that dendritic inhibition constrains excitatory recruitment and thereby regulates the magnitude of the population response during SWRs. Notably, dendritic interneurons also inhibit other inhibitory populations; accordingly, their removal led to a moderate increase in the firing of perisomatic interneurons, whereas axo-axonic activity remained largely unchanged. Thus, despite increased inhibitory recruitment, the dominant consequence of dendritic disinhibition was a substantial increase in pyramidal population activity, highlighting dendritic inhibition as a critical mechanism controlling the strength of hippocampal SWRs.

Together, these findings reveal a functional specialization of inhibitory domains during SWRs. Perisomatic inhibition acts as a gate required for the emergence of ripple oscillations, whereas dendritic inhibition fine-tunes their spectral properties by regulating the level of pyramidal recruitment. In contrast, axo-axonic inhibition contributes only modestly to SWR dynamics.

### Inhibition of inhibition emerges as an additional layer of control over SWR oscillatory dynamics

While the previous analyses revealed how distinct inhibitory populations directly regulate pyramidal cell excitability, hippocampal interneurons also form extensive reciprocal inhibitory networks. Therefore, the impact of an inhibitory population cannot be fully understood by considering only its direct synapses onto pyramidal cells, as interneurons also regulate each other and thereby indirectly control the overall inhibitory tone of the circuit. To isolate this additional level of regulation, we selectively removed interneuron-to-interneuron (IN-IN) connections while preserving all inhibitory projections onto pyramidal cells.

Specifically, we first removed the inhibitory output of perisomatic-targeting interneurons onto the remaining interneuron populations, including connections onto perisomatic, dendritic, and axo-axonic interneurons (**Figure 7a, condition 2**). We then selectively removed dendritic-targeting interneuron inhibition onto the interneuron network (**Figure 7a, condition 3**). Finally, we abolished all interneuron-to-interneuron interactions (**Figure 7a, condition 4**). Axo-axonic interneuron outputs onto other inhibitory populations were not manipulated because these cells exclusively target the axon initial segment of pyramidal neurons and, as shown previously (**Figure 6**), their contribution to SWR dynamics is minimal.

**Figure 7.**
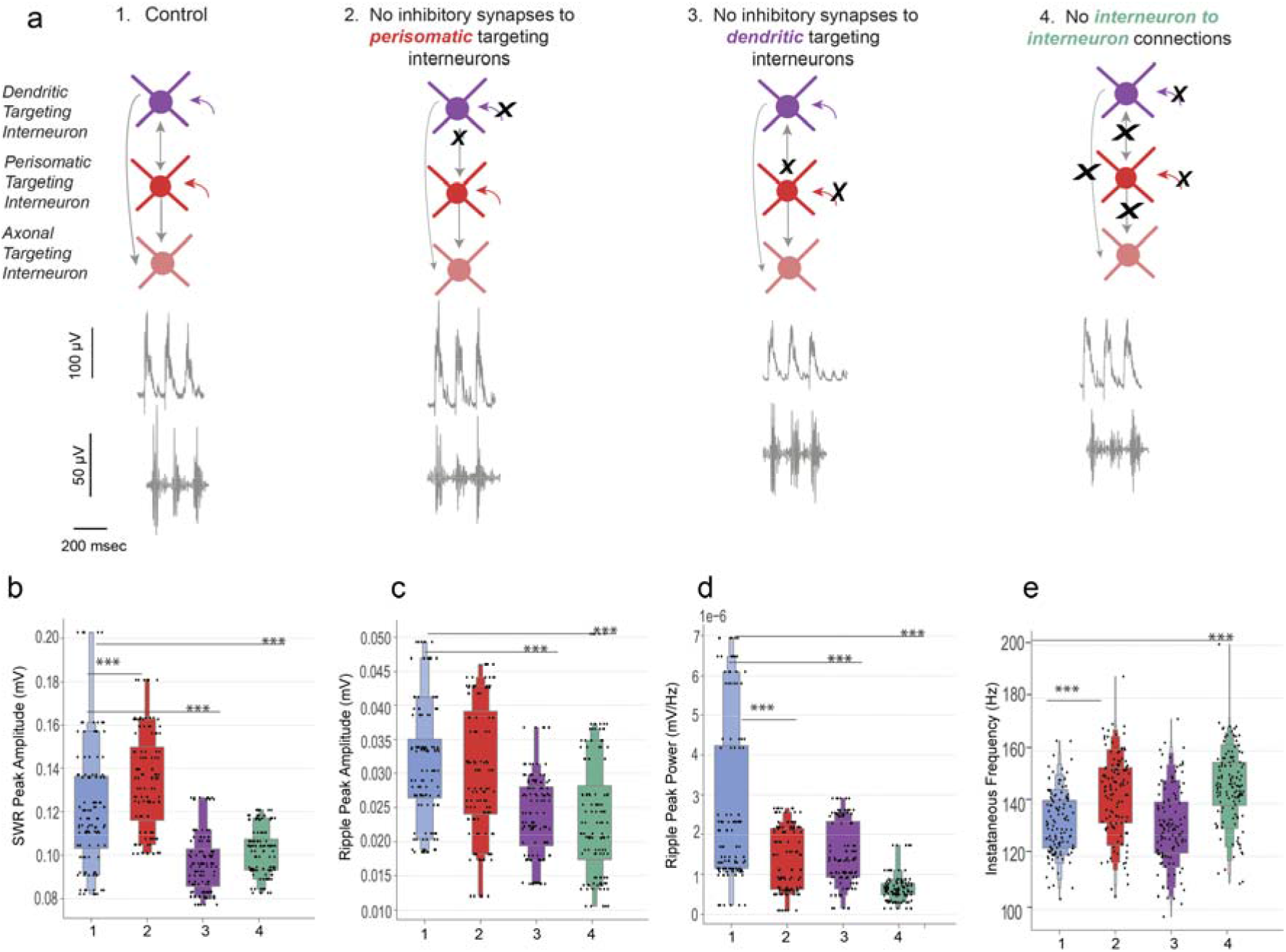
Selective disruption of interneurons connections differentially affects SWR dynamics. **(a)** Schematics of targeted in silico interneuron-to-interneuron connections disruption protocols. To isolate the role of inhibitory feedback, specific IN-IN connections were selectively severed while preserving all direct inhibitory projections onto pyramidal cells. The conditions include: Control network (1); Ablation of inhibitory synapses to perisomatic-targeting interneurons (2); Ablation of inhibitory synapses to dendritic-targeting interneurons (3); and complete ablation of all IN-IN connections across the entire network (4). Representative raw LFP sharp-wave traces (top) and band-pass filtered ripple oscillations (bottom) demonstrate that SWR generation survives these manipulations but undergoes distinct structural alterations. **(b–e)** Comprehensive quantification of SWR electrophysiological features across the reciprocal disinhibition protocols. Box plots with overlaid individual data points compare SWR raw peak amplitude (**b**), ripple peak amplitude (**c**), peak ripple power (**d**), and instantaneous ripple frequency (**e**). Disruption of local reciprocal inhibition endows a complex reorganization of the circuit’s overall inhibitory balance. For example, silencing inhibitory control over dendritic inhibitory population (Condition 3) chokes off SWR magnitude and power. Furthermore, the complete ablation of all IN-IN connections (Condition 4) dampens overall amplitude while accelerating the instantaneous ripple frequency. Thus, reciprocal interneuron connectivity acts as an independent regulatory dial essential for fine-tuning the precise spectral pacing and intensity of the fast oscillation. Data represent n = 120 analyzed SWR events per experimental condition, extracted across 10 independent, randomized simulation trials. Boxplots show mean and std values. Statistical significance was evaluated using generalized linear mixed-effects model to account for hierarchical trial variance (*p < 0.05, **p < 0.01, ***p < 0.001).

Removing perisomatic inhibition onto the interneuron network resulted in a significant increase in SWR amplitude, ripple peak amplitude, ripple power, and instantaneous ripple frequency (**Figure 7b–e**). This effect is consistent with a disinhibitory mechanism: eliminating the inhibition exerted by perisomatic interneurons onto other inhibitory populations increases the activity of these interneurons, ultimately modifying the balance of inhibition controlling pyramidal recruitment during SWRs.

In contrast, removal of dendritic inhibition from the interneuron network produced the opposite effect. SWRs exhibited reduced raw amplitude, decreased ripple peak amplitude, and lower ripple power (**Figure 7b–d**). These observations are consistent with increased activity of dendritic-targeting interneurons when released from inhibitory control. Since dendritic inhibition strongly constrains pyramidal dendritic excitation, enhanced dendritic inhibitory activity reduces pyramidal recruitment and consequently decreases the overall magnitude of SWR events. This interpretation is in agreement with our previous findings showing that complete removal of dendritic inhibition produced the opposite phenotype, namely enhanced pyramidal firing and increased SWR amplitude (**Figure 6; Supplementary Figure 6**).

Interestingly, complete removal of all interneuron-to-interneuron connections did not simply reproduce the effects of individual manipulations but revealed a complex reorganization of inhibitory balance. Global disinhibition of the interneuron network reduced SWR amplitude, ripple peak amplitude, and ripple power, indicating an overall increase in inhibitory control over pyramidal populations. At the same time, instantaneous ripple frequency was increased (**Figure 7e**), suggesting that reciprocal inhibitory interactions contribute not only to the strength of SWR events but also to the precise temporal organization of fast oscillatory activity.

Importantly, implementing the same interneuron-to-interneuron manipulations in the global inhibitory topology network produced qualitatively similar effects (**Supplementary Figure 7**). Together, these results demonstrate that interneuron-to-interneuron connectivity represents an additional regulatory layer controlling the spectral properties of SWRs. In contrast to inhibitory topology, which determines the spatial organization and coexistence of local ripple generators, local inhibitory interactions regulate the intensity and temporal structure of the oscillation itself.

### Local inhibitory topology preserves autonomous SWR generators across neighboring network subdomains

The previous analyses established that inhibitory topology determines the ability of multiple SWR generators to coexist across the hippocampal axis, whereas distinct inhibitory populations and interneuron interactions regulate the generation and spectral characteristics of individual events. However, a fundamental prediction of a locally organized inhibitory architecture remains untested: whether the failure of SWR generation in one hippocampal subdomain can influence the activity of neighboring generators.

To directly address this question, we selectively abolished perisomatic inhibition in one spatial subdomain of the network, thereby sublocally preventing SWR generation while preserving the neighboring subdomains intact (**Figure 8a**). Based on our previous simulations, we predicted that a structured inhibitory topology should maintain the independence of adjacent SWR generators, whereas a global inhibitory architecture should allow distant inhibitory interactions to influence neighboring domains.

**Figure 8.**
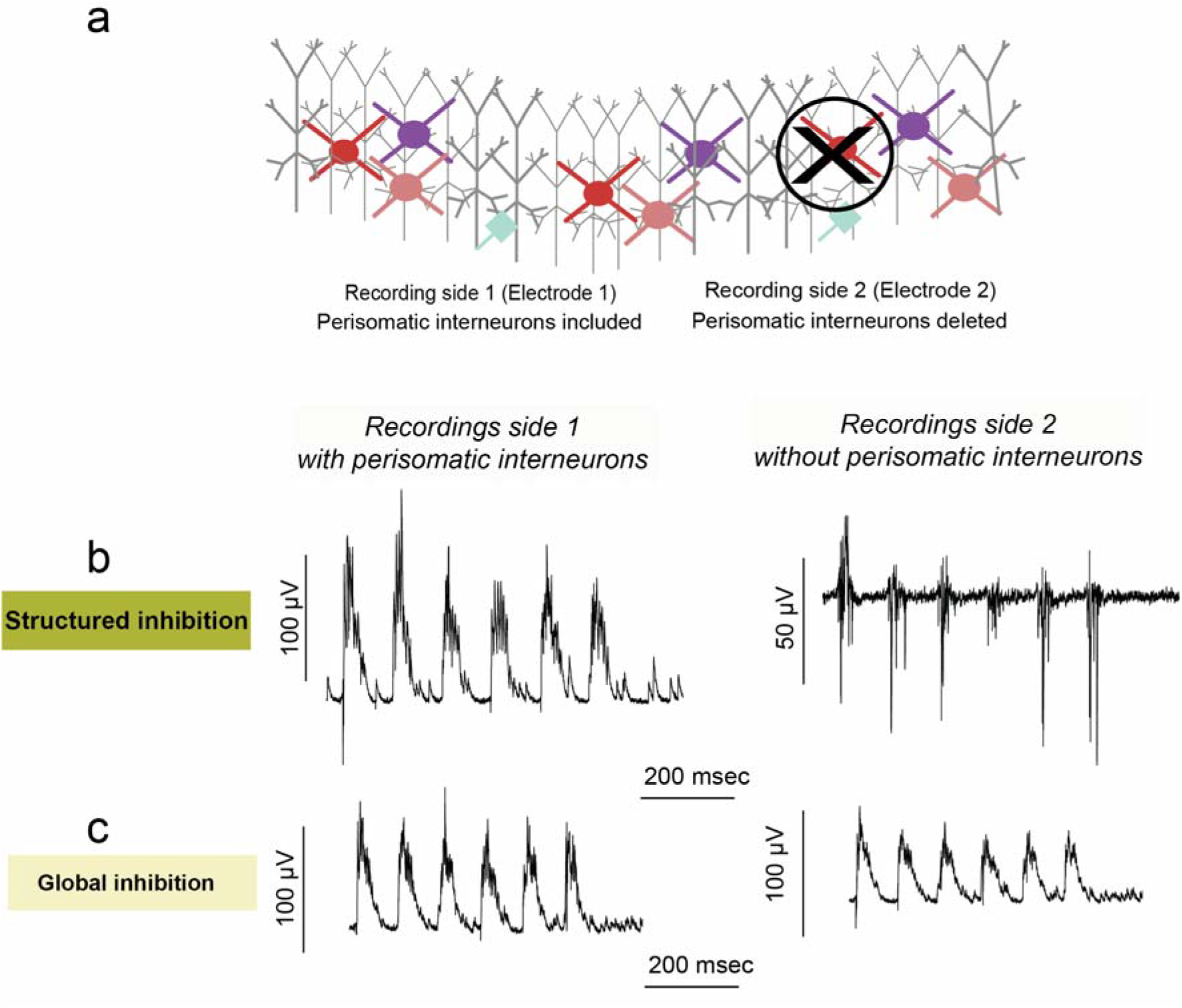
Local inhibitory topology preserves autonomous SWR generators against spatially restricted perturbations. **(a)** Schematic of the targeted in silico spatial lesion protocol. The CA1 network is divided into spatial subdomains: the left domain (Recording side 1) remains anatomically intact, while perisomatic-targeting interneurons are selectively ablated only in the right domain (Recording side 2). **(b)** Under **structured inhibition**, the localized functional failure remains restricted to the lesioned right side subdomain. The other intact subdomains maintain robust, independent SWR (left), while the lesioned subdomain exhibits a loss of organized rhythmic activity (right). This shows that distance-dependent local topology effectively isolates neighboring domains, preserving spatial autonomy. **(c)** Under **global inhibition**, the network lacks protective spatial boundaries. Long-range perisomatic projections from the intact domain (left) extend into the lesioned domain, rescuing SWR generation there (right) despite the local absence of perisomatic interneurons. Thus, we predict that structured inhibitory topology may be a critical structural requirement not only for the coexistence of multiple SWR foci, but for protecting independent memory-related computations from neighborhood failures.

Consistent with this prediction, structured inhibition preserved SWR activity in the unaffected subdomain but abolished SWRs in the subdomain without perisomatic cells. (**Figure 8b**). In sharp contrast, under global inhibitory topology, long-range perisomatic projections from the intact side propagated across the network and sustained SWR activity even in the domain lacking local perisomatic inhibition (**Figure 8c**). Thus, although global inhibitory networks are capable of generating SWRs under homogeneous conditions, they lack the ability to preserve independent local generators when the network is exposed to spatially heterogeneous perturbations.Together, these results provide evidence that structured inhibitory topology creates functionally autonomous hippocampal subdomains. Therefore, local inhibitory architecture not only allows the coexistence of multiple SWR foci but also protects individual generators from perturbations occurring in neighboring regions.

All in all, we propose that SWR generation and features are driven by inhibitory activity, but SWR spatial autonomy depends on inhibitory topology.

## Discussion

### Beyond the Clockwork: The Spatial Dimension of Inhibition

Despite being among the most synchronous events in the mammalian brain^20,59–62^, SWRs can emerge as spatially restricted events that coexist alongside neighboring ripple generators^17,22,23^. How such local autonomy is preserved within the CA1 network has remained a central paradox. Over the past decades, the field has thoroughly illuminated the “clockwork” of SWRs, firmly establishing that coordinated excitatory-inhibitory interactions, determine when neurons fire and how a rhythmic event is generated. Consequently, inhibition has been largely conceptualized as a temporal mechanism. Here, combining *in vivo* Neuropixels recordings (**Figure 2**) with an experimentally constrained biophysical model (**Figures 3-8**), we introduce an equally critical, yet underappreciated dimension of inhibition: *space*. We propose a fundamental separation between the mechanisms governing SWR *generation* and those governing SWR *spatial organization (****Figures 1,5,6,8***). While the activity of inhibitory populations acts as the temporal engine that synthesizes the ripple, the topological architecture of those connections dictates whether that activity forms a single, fragile global event or multiple, robust, independent computational assemblies.

### Distance-dependent inhibition and the emergence of autonomous domains

This topological necessity is deeply rooted in the principles of statistical physics^63^ and network theory^25^. In complex systems, small-world^63^ architectures represent an elegant thermodynamic solution to a universal challenge: how to preserve global communication while preventing runaway synchronization or temporal collapse. By combining dense local clustering with sparse long-range interactions, such networks maximize local order without sacrificing the autonomy of individual modules. Our findings suggest that the CA1 circuit may exploit analogous organizational principles. As defined by Peters’ rule and anatomical constraints, interneurons are substantially more likely to interact with neighboring cells than distant targets. By considering this spatial constraint *in silico (****Figure 3***), we revealed the function of structured inhibition (**Figure 5,8**). While a global, spatially agnostic network could technically generate an SWR event (**Figure 4**), it proved fragile under heterogeneous conditions, collapsing into competitive interactions where dominant domains suppressed weaker ones. Structured inhibition, however, acted as a spatial buffer (**Figure 5,8**). Our model captures the inherent trial-to-trial variability of SWR waveforms, such as fluctuations in spectral power and duration, which have recently been identified as signatures of underlying input diversity rather than stochastic noise. Sebastian et al. (2023) demonstrated that these reflect the specific spatial and synaptic history of the local CA1 circuit. Our findings extend this perspective by suggesting that this variability is not merely a reflection of upstream inputs, but is actively shaped and filtered by the local inhibitory architecture^18^. By partitioning the network into autonomous computational subdomains, structured inhibitory topology can ensure that each event is a local signature of the specific microcircuit it inhabits, providing a mechanistic basis for the waveform heterogeneity observed *in vivo (****Figure 2***). Thus, we propose that local inhibitory architecture prevents the network from collapsing into a single, dominant oscillation.

### Functional implications of spatial compartmentalization

The capacity to preserve multiple autonomous SWR foci has profound implications for memory processing^1–4^. During behavior and sleep, the hippocampus is bombarded by diverse, dynamically changing inputs from the CA3, entorhinal cortex, and septal nuclei^5,64^. If CA1 operated under a global inhibitory architecture, a strong input to one domain might risk hijacking the entire hippocampal axis. Instead, structured inhibition could compartmentalize the network. This modularity may provide the hippocampus with the flexibility to maintain parallel, independent memory-related computations simultaneously, without losing the capacity for large-scale synchronization and long-range propagation, when necessary.

### Segregation vs. Propagation: Stitching the Modules Together

Crucially, a sharp distinction must be drawn between the spatial *segregation* of SWRs, the focus of this study, and their *propagation*. It is well-established that SWRs can act as traveling waves across the longitudinal axis^17,23,28^. Our results in **Figure 8** show that a global inhibitory topology breaks local segregation, allowing activity to leak across the network. However, relying solely on globalized perisomatic basket cells to spread this activity leads to mutual interference and loss of computational independence, rather than organized propagation. While electrical coupling via gap junctions between PV+ perisomatic interneurons undoubtedly fortifies rapid synchronization within a local focus^65,66^, true wave propagation requires a mechanism that can safely bridge these domains. Emerging evidence highlights the existence of specialized PV+ interneuronal functional assemblies active during memory consolidation^27^. Consistent with this modular view, recent work has shown that temporally clustered SWRs, replay spatially extended trajectories and preferentially engage hippocampal-neocortical networks during consolidation^67^. The spatial compartmentalization we describe here may provide the structural substrate that allows such ripple clusters to operate independently at multiple foci, sustaining parallel replay without mutual interference: a prerequisite that the temporal clustering framework implicitly requires but does not explain. We propose that local SWR generators represent elementary computational units, and that organized traveling waves are formed by the sequential recruitment of these independent modules. We speculate that specific, long-range inhibitory projection neurons act as the connective tissue between these domains. For instance, recently identified TORO cells^68,69^ are known to propagate SWR-related information *outside* the hippocampus; an urgent and exciting open question is whether these, or analogous long-range interneurons, orchestrate the structured propagation of SWRs *within* the CA1 axis.

### A Hierarchy of Control: Gating, Sculpting, and Disinhibition

Beyond spatial boundaries, our study reveals a strict hierarchical division of labor among inhibitory cell types. Although PV+ interneurons are frequently treated as a monolithic population, our targeted deletions demonstrate that their subcellular targets define distinct computational roles (**Figure 6-7**). Fast perisomatic inhibition acts as a binary gate; without it, the SWR rhythm cannot emerge. In contrast, dendritic inhibition does not dictate *if* an SWR exists, but rather *how loudly* it is expressed. Dendritic disinhibition triggered massive pyramidal recruitment, severely inflating SWR amplitude, duration, and ripple power (**Figure 6, Supplementary Figure 6**). Thus, dendritic inhibition functions as a continuous gain-control mechanism, restricting the size of the participating excitatory ensemble and preventing pathological runaway excitation. Furthermore, reciprocal interneuron-to-interneuron (IN-IN) disinhibition constitutes a third regulatory layer (**Figure 7**). These local inhibitory feedback loops do not merely modulate amplitude but are essential for fine-tuning the precise spectral pacing of the fast oscillation.

### Biological Fidelity over Computational Complexity

The design of our *in silico* network reflects deliberate biological constraints rather than computational limitations. We exclusively modeled inhibitory populations because they are the undisputed drivers of the SWR rhythm. While the CA1 network features a vast array of other inhibitory classes, most notably somatostatin (SOM+) and vasoactive intestinal peptide (VIP+) interneurons, extensive experimental evidence demonstrates that these populations are actively silenced during CA1 SWRs^9,55,56,70^. Their deliberate exclusion thus preserves the strict biological fidelity of the physiological ripple state. Similarly, we deliberately restricted our topological manipulations to the inhibitory network, maintaining a constant, biologically structured excitatory (pyramidal) topology across all simulations. While certain theoretical frameworks emphasize excitatory interactions in SWRs^14,31,71^, pyramidal cells exhibit characteristically sparse firing rates and low overall participation probabilities during biological hippocampal SWRs^72^. This sparse excitatory activity is poorly suited to rapidly enforce rigid spatial boundaries across the network. Instead, the dense, high-frequency bursting of interneurons uniquely positions the inhibitory network to contain and compartmentalize runaway synchronization. Furthermore, we deliberately restricted our model to chemical synaptic transmission, omitting the electrical coupling (gap junctions) known to tightly synchronize especially PV+ interneurons^65,73^. While gap junctions are undeniably critical for orchestrating micro-second interneuronal synchrony during biological SWRs, their connections are anatomically restricted to immediate, local neighbors. By forcing our model to rely entirely on chemical synapses, we subjected our spatial segregation hypothesis to a much more rigorous topological test. Because gap junctions selectively bind adjacent cells, their inclusion *in vivo* would theoretically act as a localized amplifier, further fortifying the “small-world” clustering and structural robustness of the local compartments we observed here. Finally, we purposefully isolated the CA1 network from external, extra-hippocampal afferents, including the entorhinal cortex (EC) and medial septum. This isolation was a strategic necessity: to test whether SWR spatial segregation is an *intrinsic* architectural property of the CA1, we had to test if it emerges from local topology alone, rather than being inherited from patterned external commands. However, the absence of these external inputs remains a critical boundary of our current model. *In vivo*, the hippocampus is continuously bombarded by diverse, competing streams of information^74^. We hypothesize that intrinsic compartmentalization may exists precisely to manage this immense cognitive load, providing the hippocampus with the structural flexibility to process multiple, parallel memory traces simultaneously without interference. Future studies mapping these extra-hippocampal inputs onto our blueprint will likely reveal how the “walls” of these spatial compartments are dynamically erected, dissolved, or shifted *in vivo* to route specific memories.

## Conclusion

Together, our findings highlight a previously unrecognized separation between the mechanisms that generate hippocampal SWRs and those that organize them across space. We propose a unifying conceptual framework: inhibitory *activity* determines whether a ripple emerges and defines its spectral identity, while inhibitory *topology* dictates whether multiple ripple generators can coexist as autonomous units. Inhibition may no longer be viewed solely as the temporal conductor of hippocampal rhythms, but equally as the spatial architect that dictates the boundaries, autonomy, and integration of memory traces.

## Author contributions

Conceptualization, Original Hypothesis and Design: AT

Interpretation of the Results: AT with inputs from DP

Computational Modeling: AT

Analysis: AT, DP, AK, RF

Manuscript Preparation: AT

Manuscript editing: AT, DP, AK

Project administration: AT Resources: DS

Funding: DS, AT

## Supporting information

Supplementary Infortmation

## Acknowledgements

AT is supported by the Schering Stiftung Fellowship, and by the NeuroCure Exc-2049, Postdoc Fellowship. D.S. lab is supported by the Einstein Foundation Berlin, the European Research Council (ERC) under the Europeans Union’s Horizon 2020 research and innovation program (BrainPlay grant, agreement no. 810580), the German Research Foundation (Deutsche Forschungsgemeinschaft [DFG], SFB-958 – project 184695641, project 431572356, FOR 3004 – project 415914819, SFB 1315 – project 327654276 and under Germany’s Excellence Strategy – Exc-2049-390688087 NeuroCure), and the Federal Ministry of Education and Research (BMBF, SmartAge – project 01GQ1420B). We thank very much John Tukker and Jeremie Sibille for discussions about the experimental results, Spiros Chavlis for advising on the model network implementation and Nikolaus Maier for feedback on the manuscript.

## Code and Data availability

All codes/scripts and datasets required to reproduce the results and figures, as well as all statistical analyses, are accessible in the ModelDB database. They are also available on GitHub: https://github.com/AlexandraTzilivaki/Tzilivakietal2026

For questions contact

## Online Methods

### 1.0 Experimental Dataset analysis

The *in vivo* Neuropixels recordings analyzed here were collected by Kala et al. (2026)² , and are described in full detail in that study. Briefly, tangential Neuropixels 1.0 probe insertions were performed in 14 adult male C57BL/6J mice under head-fixed conditions, approved by the Landesamt für Gesundheit und Soziales Berlin (G0298/18) and conducted according to institutional and national guidelines for animal welfare. No new animal experiments, surgeries, or recordings were performed for the present study; we received the previously acquired electrophysiological dataset for reanalysis.

### Detection and spatiotemporal analysis of *in vivo* SWRs

LFP signals obtained from Neuropixels recordings were first corrected for the temporal offsets introduced by the multiplexed analog-to-digital converters (ADCs) of the Neuropixels 1.0 probe. Because channels are sequentially sampled within each ADC, this results in cumulative inter-channel delays of up to approximately 0.4 ms. These offsets were corrected before subsequent analyses to ensure accurate temporal comparison of ripple activity across recording sites. For ripple detection, LFP signals were band-pass filtered between 110 and 240 Hz using a fourth-order Butterworth filter. The analytic signal was obtained using the Hilbert transform, and ripple power was defined as the absolute value of the transformed signal. Candidate ripple events were identified when the ripple envelope exceeded a threshold of five standard deviations above baseline. Events with durations shorter than 50 ms or longer than 200 ms were excluded from further analyses. The spatial organization of SWR events was examined by comparing ripple peak times across recording channels spanning up to approximately 1.3 mm along the longitudinal CA1 axis. To quantify the relationship between spatial location and temporal occurrence of ripple events, Pearson’s correlation analysis was performed between the anatomical position of each recording channel and the corresponding ripple peak time. To further examine the degree of synchrony and potential spatiotemporal organization of SWR activity across the longitudinal axis, cross-correlograms of ripple peak times between recording channels were calculated in MATLAB (MathWorks).

### 2.1 Model implementation and availability

Simulations were performed with NEURON (v7.6)^75^on a High-Performance Computing Cluster, utilizing 111 CPU cores on a 64-bit CentOS Linux operating system. All codes/scripts and datasets required to reproduce the results and figures, are accessible in the ModelDB database and github.com/AlexandraTzilivaki.

### 2.2 Neuronal populations

#### Single-cell biophysical models and validation

Individual neuronal populations were represented by multicompartment conductance-based models implemented in NEURON using Hodgkin–Huxley type ionic mechanisms as per^30^. The level of morphological complexity was adapted to the anatomical and electrophysiological characteristics of each neuronal class. CA1 pyramidal neurons were represented by a detailed multicompartment model containing somatic, axonal, apical, and basal dendritic compartments, enabling realistic spatial segregation of excitatory and inhibitory synaptic inputs. In contrast, inhibitory interneurons were represented by simplified multicompartment models capturing their characteristic fast-spiking electrophysiological behavior while maintaining their distinct synaptic targeting domains. The intrinsic electrophysiological properties of each neuronal type were validated against experimental measurements^32–34,38,39,41,42,45,46,51,54^. Passive and active membrane properties, including input resistance , and rheobase current, were adjusted to reproduce the characteristic physiological behavior of CA1 pyramidal neurons and perisomatic-, dendritic-, and axonal-targeting interneurons (**Supplementary Tables 1-3**).

#### CA1 pyramidal neurons

The CA1 pyramidal cell model was adapted from the multicompartment model of ^30,38^ The model consisted of a somatic compartment, an axon, apical dendritic compartments representing proximal, medial, and distal regions of the apical trunk, and basal dendritic compartments reproducing the basal dendritic tree. The model included active conductances responsible for sodium, potassium, calcium, and hyperpolarization-activated currents, as well as calcium buffering and calcium-dependent potassium mechanisms required for realistic action potential generation, dendritic excitability, and afterhyperpolarization dynamics. The spatial organization of the model was preserved within the CA1 architecture, with the soma positioned in the stratum pyramidale, basal dendrites extending into stratum oriens, and apical dendrites extending through stratum radiatum toward stratum lacunosum-moleculare. CA1 pyramidal neurons received excitatory Schaffer collateral inputs from CA3 onto their apical dendritic compartments, while inhibitory inputs were spatially segregated according to their target domains: perisomatic inhibition targeted the soma and proximal dendrites, dendritic inhibition targeted the apical and basal dendritic compartments, and axonal inhibition targeted the axon initial segment. The maximum conductances and distribution of ionic mechanisms are provided in **Supplementary Tables 1-4**.

#### Inhibitory interneuron models

Perisomatic-, dendritic-, and axonal-targeting interneurons were modeled as multicompartment conductance-based neurons with simplified morphologies while preserving their characteristic intrinsic electrophysiological profiles and synaptic target specificity^32,38–40^. Perisomatic- and axonal-targeting interneurons consisted of 17 compartments, whereas dendritic-targeting interneurons consisted of 13 compartments. Interneuronal models included sodium, delayed rectifier potassium, A-type potassium, calcium, and calcium-dependent potassium conductances necessary to reproduce their firing dynamics. All inhibitory populations received feedforward excitatory input from CA3 Schaffer collaterals and feedback excitation from CA1 pyramidal neurons. Perisomatic- and dendritic-targeting interneurons additionally formed reciprocal inhibitory interactions within the inhibitory network, whereas axonal-targeting interneurons selectively targeted pyramidal neurons and did not inhibit other interneuron populations. The distribution and maximal conductances of all ionic mechanisms are summarized in **Supplementary Table 2**. <u>Synaptic mechanisms</u>

The CA1 microcircuit included both glutamatergic and GABAergic synaptic transmission. Excitatory synapses were mediated by AMPA and NMDA receptors, with NMDA receptor-mediated currents restricted to pyramidal neurons, whereas interneurons received excitatory input through AMPA receptors. Inhibitory transmission consisted of both GABAa and GABAb-mediated synapses. Synaptic conductances were modeled as double exponential functions characterized by their rise and decay kinetics. Synaptic parameters, including maximal conductance and time constants, were adapted from experimentally constrained CA1 network models^30,38^ and are provided in **Supplementary Table 3**. To validate synaptic efficacy, the amplitude and kinetics of excitatory and inhibitory postsynaptic currents (EPSCs and IPSCs) generated by each synaptic pathway were evaluated using voltage-clamp simulations in NEURON and compared with experimentally reported values.

### 2.4 Connectivity

The CA1 network consisted of 150 pyramidal cells and 18 inhibitory interneurons, preserving an approximate excitatory-to-inhibitory ratio of 10:1 consistent with the cellular composition of the hippocampal CA1 region. The inhibitory population was further subdivided into 9 perisomatic-targeting interneurons (basket cells), 6 dendritic-targeting interneurons (bistratified cells), and 3 axonal-targeting interneurons (axo-axonic cells). Interneurons were spatially distributed along the longitudinal axis of the CA1 network to ensure representation of all major inhibitory domains across the circuit and to allow an unbiased investigation of how structured inhibitory topology contributes to SWR generation and spatial segregation. Excitatory drive to the CA1 network was provided by artificial CA3 Schaffer collateral afferents. Based on experimental estimates of hippocampal connectivity, pyramidal cells received a higher convergence of CA3 inputs compared with inhibitory populations, reflecting the strong excitatory drive of Schaffer collateral projections onto CA1 pyramidal neurons^33,41,76^. CA3 inputs targeted the apical and basal dendritic compartments of pyramidal cells through AMPA- and NMDA-mediated synapses, whereas inhibitory populations received a lower degree of feedforward excitatory input. Within the CA1 circuit, pyramidal cells formed sparse excitatory connections with other pyramidal cells and provided excitatory feedback to all inhibitory populations. Inhibitory connectivity was organized according to distinct subcellular targeting domains. Perisomatic-targeting interneurons innervated the soma and proximal dendrites of pyramidal cells, dendritic-targeting interneurons targeted the apical and basal dendritic arbor, and axonal-targeting interneurons selectively innervated the axon initial segment, thereby regulating distinct aspects of pyramidal cell integration and action potential generation. In addition to their inhibitory control of pyramidal neurons, inhibitory populations formed a structured interneuron network. Perisomatic-targeting interneurons established reciprocal inhibitory connections with other perisomatic-targeting interneurons and provided inhibition onto dendritic-targeting and axonal-targeting interneurons. Likewise, dendritic-targeting interneurons inhibited all interneuron populations, including perisomatic-, dendritic-, and axonal-targeting cells. In contrast, axonal-targeting interneurons selectively targeted pyramidal neurons and did not provide inhibitory connections onto other interneuron classes. The number, spatial organization, and target specificity of synaptic connections were constrained by experimental anatomical and physiological data from the hippocampal CA1 microcircuit^32–34,38,40,55^. This architecture enabled the investigation of how the spatial organization and interactions between distinct inhibitory domains regulate the emergence and segregation of memory-related SWR activity.

### 2.5 Three-dimensional organisation of the CA1 network

To reproduce the anatomical organization of the hippocampal CA1 region, neurons were positioned within a three-dimensional spatial framework approximating the characteristic curved “banana-shaped” geometry of the CA1 pyramidal cell layer. The longitudinal axis of CA1 was represented along the x-axis, whereas the y-axis represented the laminar organization perpendicular to the pyramidal cell layer, and the z-axis represented the transverse dimension of the tissue. Pyramidal cell somata were distributed within the stratum pyramidale following a curved spatial arrangement generated by a parabolic function along the longitudinal axis, thereby reproducing the characteristic arc-shaped morphology of the CA1 pyramidal layer. A total of 150 pyramidal cells were positioned on the longitudinal axis, with an average intersomatic distance of approximately 20–25 μm, consistent with the dense packing of CA1 pyramidal neurons. Their somatic positions were restricted to a ∼100 μm thick pyramidal layer. Inhibitory interneurons were placed according to their known laminar distribution and target domains. Perisomatic-targeting interneurons and a proportion of dendritic- and axonal-targeting interneurons were positioned within or close to the stratum pyramidale, reflecting their anatomical localization around the pyramidal cell layer. Their positions were distributed along the longitudinal axis to provide inhibitory coverage throughout the network. The z-axis coordinates of all neurons were randomly assigned within a narrow range, generating a three-dimensional network volume while preserving the overall laminar organization. This spatial organization allowed the implementation of distance-dependent connectivity rules, where synaptic connections could be constrained by the physical proximity of pre- and postsynaptic neurons. The resulting three-dimensional CA1 architecture enabled the investigation of how spatially organized inhibitory connectivity contributes to the generation, localization, and propagation of sharp-wave ripple activity.

### 2.5 Distance dependent rule, Structured inhibitory topology

Synaptic connectivity in the structured CA1 network was constrained by the spatial organization of neurons along the longitudinal axis of CA1 as per ^26^. This approach was motivated by experimental anatomical data demonstrating that hippocampal connectivity is spatially organized and limited by the overlap between presynaptic axonal projections and postsynaptic target domains, consistent with the principles of Peters’ rule. Given that CA1 axonal arbors typically span several hundred micrometers, candidate presynaptic partners were restricted to a biologically realistic interaction range of approximately 600–900 μm. For each postsynaptic neuron, presynaptic partners were randomly selected from the appropriate source population. The distance (d) along the longitudinal axis was calculated as:

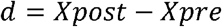

If a selected presynaptic neuron was located outside the permitted spatial range, a new candidate was randomly sampled until a valid connection within the distance constraint was identified. Inhibitory connections were restricted to neurons located within approximately 905 μm, whereas pyramidal cell-mediated excitatory connections were restricted to neurons within approximately 800 μm. Self-connections were excluded. To account for variability in the exact anatomical arrangement of local microcircuits, each simulation trial was generated using a different random realization of the connectivity matrix. Thus, although all trials preserved the same distance-dependent connectivity rules, cell-type-specific targeting, and overall connection statistics, the exact identities of connected pre- and postsynaptic neuronal pairs varied across trials. For each established synaptic connection, the synaptic strength was scaled according to a Gaussian function of intersomatic distance:

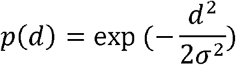

*With* σ*=600* μ*m*

The final synaptic weight was then calculated as:

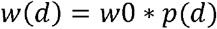

Where w0 is the baseline synaptic weight and p(d) is the distance-dependent scaling factor. Thus, nearby neurons formed stronger effective synaptic connections, whereas more distant neurons within the allowed axonal range formed weaker connections. Neurons beyond the spatial cutoff were not connected.

### Distance Dependent rule at the global inhibitory topology network

To investigate the role of spatially organized inhibition in SWR generation and propagation, we generated an alternative CA1 network configuration in which inhibitory connectivity was randomized across the septotemporal axis. In this model, inhibitory interneuron-to-pyramidal cell and interneuron-to-interneuron connections were established independently of intersomatic distance, thereby abolishing the local topological organization of inhibitory interactions. Presynaptic inhibitory partners were randomly selected from the corresponding interneuron population while preserving the same convergence, synaptic target domains, and baseline synaptic parameters as in the structured network. Self-connections were excluded. Importantly, the spatial organization of the excitatory network was maintained. Sparse pyramidal cell-to-pyramidal cell connections and pyramidal cell excitation onto interneurons continued to follow the distance-dependent connectivity rule. Thus, the global inhibitory topology model selectively disrupted the spatial organization of inhibitory pathways while preserving excitatory distance-dependent connectivity and the subcellular target specificity of the different interneuron classes. This manipulation allowed us to directly test whether local inhibitory topology is required for the generation, localization, and propagation of SWR activity.

### 2.6 Input

External excitatory drive to the CA1 network was provided by a population of artificial CA3 Schaffer collateral afferents. They were modeled as Poisson spike generators, producing stochastic spike trains with a predefined mean firing rate. For each simulation trial, a different random realization of the presynaptic activity was generated by changing the random seed while preserving identical input statistics across simulations. To introduce temporally structured oscillatory activity, the generated Poisson spike trains were filtered according to a sinusoidal probability function:

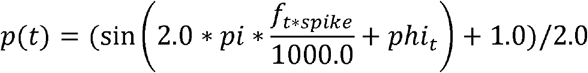

where t corresponds to the spike time in milliseconds, f represents the frequency of the oscillatory modulation, and represents the phase offset in radians.

For each generated spike, the probability P(t) was evaluated and spikes were preferentially retained during high-probability phases of the oscillatory cycle. Specifically, spikes were accepted when P(t)>0.7 and an additional random number satisfied r<P(t)/2, resulting in stochastic but phase-biased oscillatory input patterns. The random component of the selection process ensured that input spike trains remained irregular rather than forming a perfectly periodic pattern. An additional noise parameter allowed spikes to be accepted independently of the oscillatory modulation, enabling control over the level of temporal randomness. The final accepted spike times were temporally shifted according to the specified input delay and delivered to the CA1 network through Schaffer collateral synapses. Spikes occurring outside the simulation time window were discarded. This approach generated trial-to-trial variability in presynaptic activity while maintaining a controlled oscillatory temporal structure.

In addition to the structured oscillatory drive, neurons received stochastic background synaptic input to mimic the ongoing fluctuations present in biological neural circuits. This background activity introduced variability in the membrane potential and network dynamics, preventing unrealistically deterministic responses to identical inputs and increasing the robustness of SWR generation across independent simulation trials.

### Robustness/sensitivity analysis

To evaluate the robustness of the CA1 network model and ensure that the emergence of SWR activity was not dependent on a narrowly tuned parameter set, we performed a sensitivity analysis by systematically perturbing key synaptic parameters. Specifically, the strength of external CA3 Schaffer collateral inputs and the synaptic strengths of the intrinsic CA1 connectivity pathways were independently increased or decreased by 20–30% relative to their baseline values. For each perturbed condition, the resulting network activity was analyzed using the same SWR detection and characterization pipeline applied to the baseline simulations, including measurements of SWR amplitude, duration, instantaneous frequency, ripple-band power, and peak ripple frequency. Despite these substantial variations in excitatory drive and recurrent synaptic efficacy, the network consistently generated physiologically realistic SWR events with preserved temporal and spectral characteristics. These analyses demonstrate that SWR generation in the model does not rely on precise fine-tuning of individual parameters, but rather emerges robustly from the experimentally constrained cellular properties, synaptic organization, and connectivity rules incorporated into the CA1 microcircuit.

### Neuronal population participation during SWRs

Consistent with experimental observations showing that only a sparse subset of CA1 pyramidal neurons participates in individual SWR events^46^, the model was constrained such that approximately 10-25% of the pyramidal population was recruited during each SWR. Within the detected SWR time window, neuronal activity was quantified separately for pyramidal cells, perisomatic-targeting interneurons, dendritic-targeting interneurons, and axonal-targeting interneurons.

For each SWR event, the number of active neurons in each population was determined as the number of cells generating at least one action potential during the event. The participation ratio was calculated as the fraction of active neurons relative to the total number of neurons within the corresponding population:

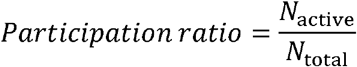

where *N*_active_ represents the number of neurons that fired during the SWR and *N*_total_ represents the total number of neurons in that population. To estimate the firing activity of participating neurons, the number of action potentials generated by each active neuron within the SWR window was calculated, and the mean number of spikes per active cell was determined for each population and each individual SWR event. This approach allowed us to quantify both the proportion of recruited neurons and the firing intensity of the participating population across different simulation conditions and independent network realizations

### 2.7 Simulation protocols

The network model was simulated for approximately 15,000 milliseconds (ms) with a time step of 0.1 ms.

### 2.8 Simulation of multi electrode recordings

To enable direct comparison between simulated and experimental recordings, in silico multi-electrode local field potential (LFP) recordings were performed using virtual extracellular electrodes implemented in the NEURON simulation environment following the extracellular recording approach described by ^71,77^. The extracellular potential was calculated from the summed transmembrane currents generated by the network and recorded at defined spatial locations around the CA1 microcircuit. Virtual recording electrodes were positioned within the stratum pyramidale, close to the somatic layer of CA1 pyramidal cells, matching the anatomical location of the in vivo recordings. To reproduce the spatial arrangement of the experimental Neuropixels recordings, electrode positions were maintained at the same laminar and transverse coordinates (y- and z-axes), while the recording sites were separated along the longitudinal CA1 axis (x-axis). Two recording sites, corresponding to septal and temporal CA1 regions, were placed approximately 1.4–1.5 mm apart, reproducing the spatial separation between the recording channels used in vivo. The virtual LFP signals were sampled at 10 kHz and recorded throughout all simulation protocols and independent network realizations. This recording configuration enabled the direct characterization of the spatiotemporal organization of simulated SWR events and allowed us to determine whether SWRs emerged as spatially localized, multifocal events or as synchronized oscillations propagating across the longitudinal CA1 axis.

### 2.8 Detection and Spectral analysis of SWRs

Simulated sharp-wave ripple (SWR) events were detected from model-generated virtual local field potential (LFP) recordings using an analysis pipeline analogous to those commonly applied to experimental electrophysiological recordings, as we did in ^53^. Virtual LFP signals were recorded from in silico electrodes positioned at septal and temporal locations and sampled at a frequency of 10 kHz. For each simulation condition, LFP recordings from 10 independent simulation trials were analyzed. To avoid initialization artifacts associated with the onset of network activity, the first 300 ms of each recording were excluded. The LFP signal was baseline-corrected by subtracting the 25th percentile of the trace, and time was expressed in milliseconds. SWR candidate events were identified from a smoothed LFP trace obtained using a 100-point moving average filter (corresponding to approximately 10 ms). Positive LFP deflections were detected using a peak-finding algorithm, and only events with peak amplitudes exceeding 0.05 mV (50 μV) were considered as candidate SWRs. The onset and offset of each event were determined using a lower amplitude threshold of 0.005 mV (5 μV), corresponding to the points at which the LFP crossed this threshold. Candidate events with durations shorter than 20 ms or longer than 500 ms were discarded. In addition, peaks separated by less than 30 ms were not considered independent events, preventing multiple detections of the same SWR event. For each accepted SWR event, both temporal and spectral features were quantified. SWR amplitude was defined as the peak amplitude of the smoothed LFP signal, while SWR duration was calculated as the time interval between event onset and offset. Instantaneous ripple frequency was estimated from the ripple-band component of the LFP obtained using a sixth-order zero-phase Butterworth band-pass filter between 80 and 250 Hz. The analytic signal was calculated using the Hilbert transform, and instantaneous frequency was derived from the temporal derivative of the unwrapped phase. The mean instantaneous frequency within the detected SWR window was used as the event’s instantaneous ripple frequency. To quantify ripple spectral properties, the LFP segment corresponding to each detected SWR was filtered in the 80–200 Hz frequency range using a fifth-order zero-phase Butterworth filter. Power spectral density was estimated using Welch’s method, and the maximum power value was defined as the peak ripple power, whereas the frequency at which this maximum occurred was defined as the peak ripple frequency. Additionally, the maximum amplitude of the ripple-band filtered signal was quantified as the peak ripple amplitude, and the maximum amplitude of the unfiltered LFP trace within the same SWR window was quantified as the peak raw LFP amplitude.

### 2.9 Statistical analysis

#### Generalized linear mixed-effects model

Statistical analyses were performed in R using generalized linear mixed-effects models implemented in the *lme4* package. To account for the hierarchical organization of the simulated data, with multiple SWR events generated within individual simulation trials, trial identity and ripple identity were included as random intercept effects. Continuous SWR properties, including amplitude, duration, instantaneous frequency, peak power, peak frequency, peak ripple amplitude, and peak raw signal amplitude, were analyzed using generalized linear mixed-effects models with a Gamma error distribution and a log link function. Fixed effects included simulation model, recording location (electrode 1 versus electrode 2), their interaction, and ripple order within each trial (centered ripple number) when appropriate. The significance of model effects and recording location effects was assessed by likelihood ratio tests comparing the full model against reduced models lacking the corresponding fixed effects or against a null model containing only random effects. SWR occurrence probability was analyzed using a generalized linear mixed-effects model with a binomial distribution and logit link function, including the same fixed and random effect structure. Post hoc comparisons between predefined simulation conditions and between recording locations were performed using estimated marginal means (*emmeans* package). P values were corrected for multiple comparisons using the Benjamini–Hochberg false discovery rate (FDR) procedure. Satistical analysis shown in figure 2 was performed using the upaired student’s Ttest. Statistical analysis shown in Supplementary Figure 6, was performed using the Mann Whitney test for datasets with non uniform distributions. A significance threshold of *P* < 0.05 was used for all analyses.

