## Supplementary Infortmation for "Local inhibitory topology dictates the spatial compartmentalization of hippocampal sharp-wave ripples"

**Supplementary Material**

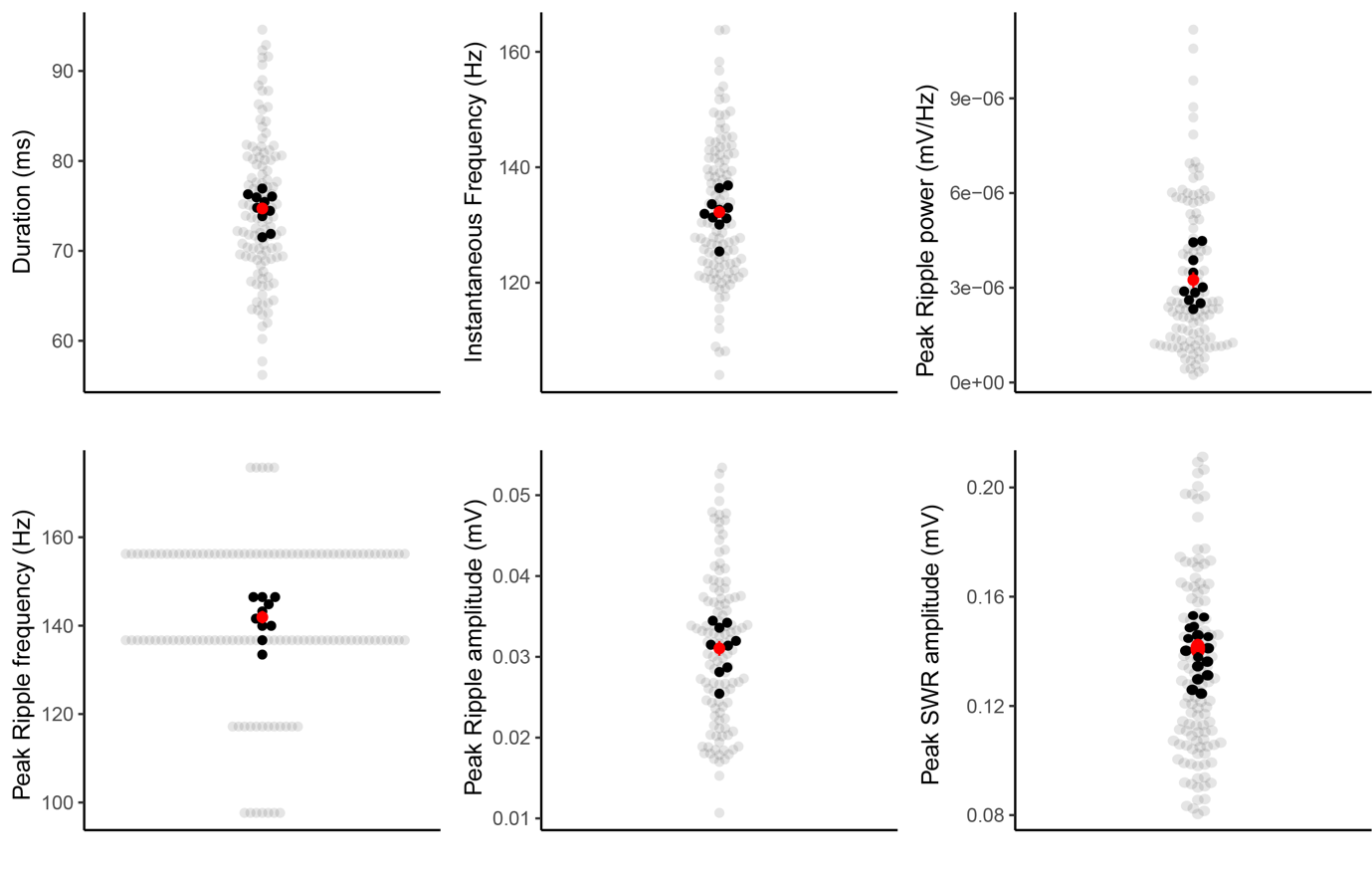
**Supplementary Figure 1 (related to Figure 3) Physiological variability of SWR dynamics in the network model.** 120 individual SWR events extracted from 10 independent simulation trials across both electrodes, demonstrating trial-to-trial inherent variability of SWR dynamics, including SWR peak amplitude, SWR duration, ripple peak power, and instantaneous ripple frequency. The observed variability in spectral power and frequency distributions matches the ranges reported in in vivo hippocampal recordings (Sebastian et al., 2023), confirming that the model reproduces the stochastic nature of biological SWRs.

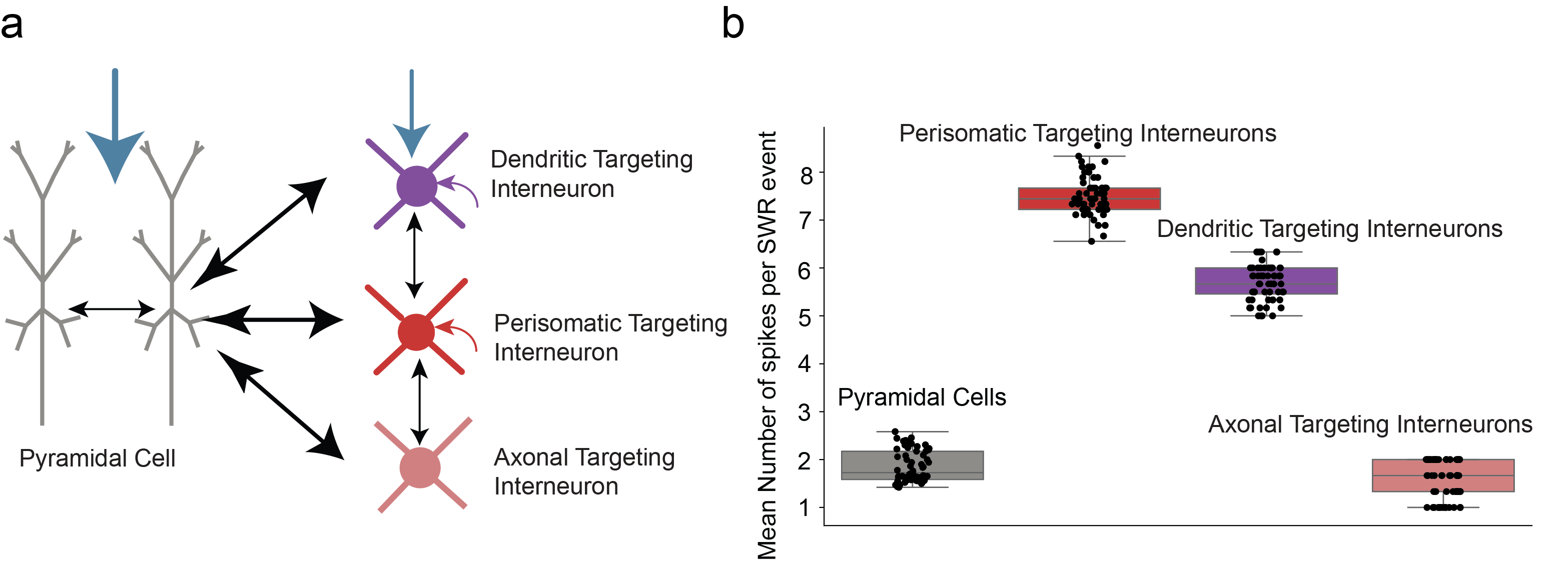

**Supplementary Figure 2 (related to Figure 3) Neuronal participation and synaptic input strength during in silico SWR events.** **(a)** Schematic representation of the CA1 microcircuit connectivity and input architecture. Arrow thickness indicates the relative strength of excitatory synaptic drive onto each neuronal population. Pyramidal cells receive the strongest excitatory drive (thicker blue arrow), reflecting experimental estimates of Schaffer collateral input dominance, while inhibitory populations receive relatively weaker, feedforward excitatory drive. Bidirectional arrows between pyramidal cells and inhibitory interneurons indicate reciprocal excitatory-inhibitory coupling. **(b)** Quantification of neuronal firing activity during individual SWR events. Box plots with overlaid individual data points show the mean number of spikes per event for pyramidal cells, perisomatic-targeting interneurons, dendritic-targeting interneurons, and axonal-targeting interneurons. These spiking activity was rigorously constrained by established experimental data.

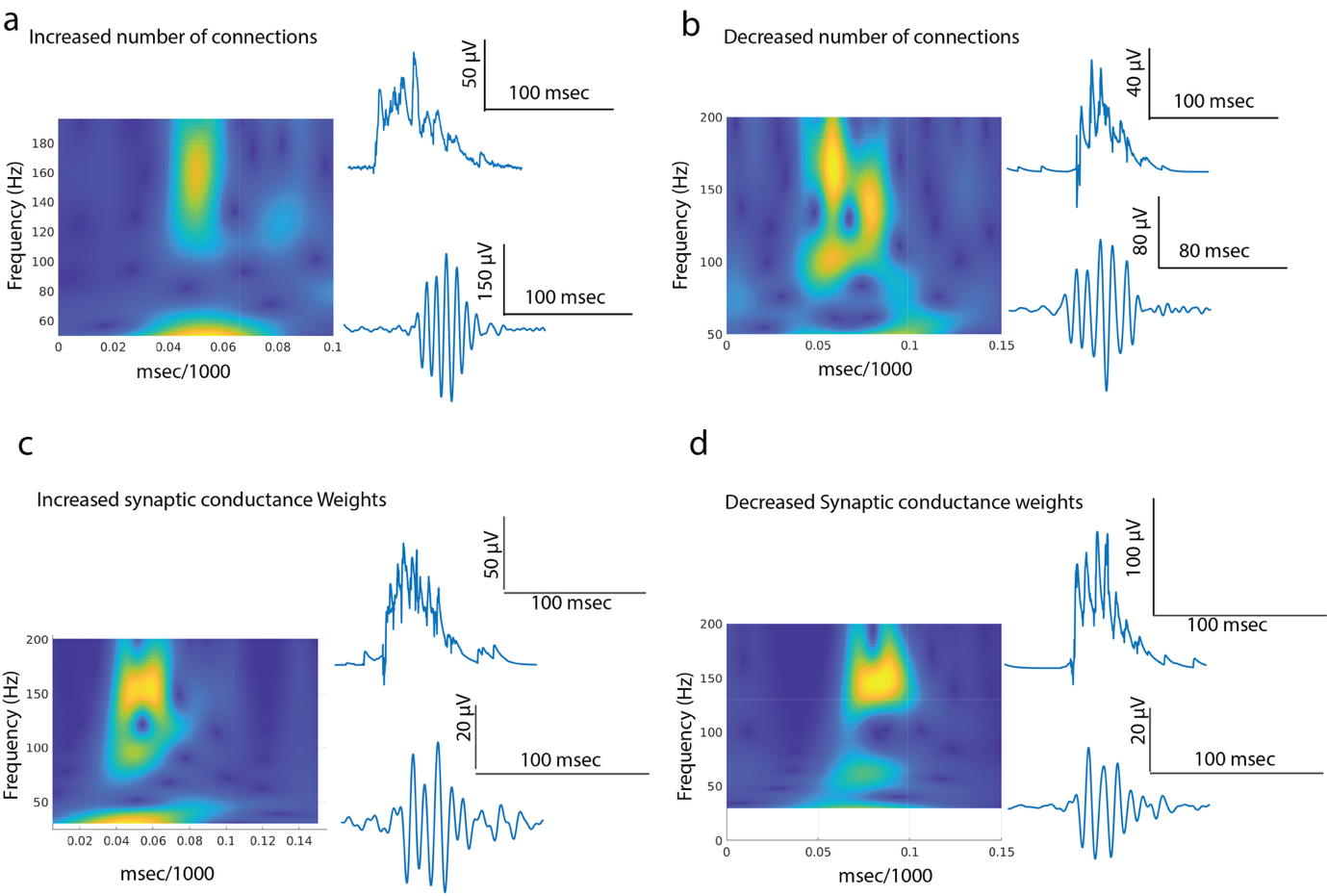

**Supplementary Figure 3 (related to Figure 3) Robustness and sensitivity analysis of SWR generation in the CA1 network model.** To ensure that the emergence of SWR activity is a robust property of the biologically constrained network architecture rather than the result of narrowly tuned parameters, we performed a systematic sensitivity analysis. We independently perturbed the strength of external CA3 Schaffer collateral excitatory drive and the synaptic conductances of intrinsic CA1 pathways by ±20–30% relative to baseline values. **(a, b)** Effect of increasing or decreasing the number of synaptic connections (variations in connectivity density). **(c, d)** Effect of increasing or decreasing synaptic conductance weights for recurrent connections. For each condition, representative time-frequency spectrograms (left) and corresponding raw LFP and band-pass filtered ripple traces (right) demonstrate that SWR activity remains stable and physiologically characteristic across all perturbations. These results confirm that SWR dynamics in the model emerge robustly from the experimentally constrained cellular properties.

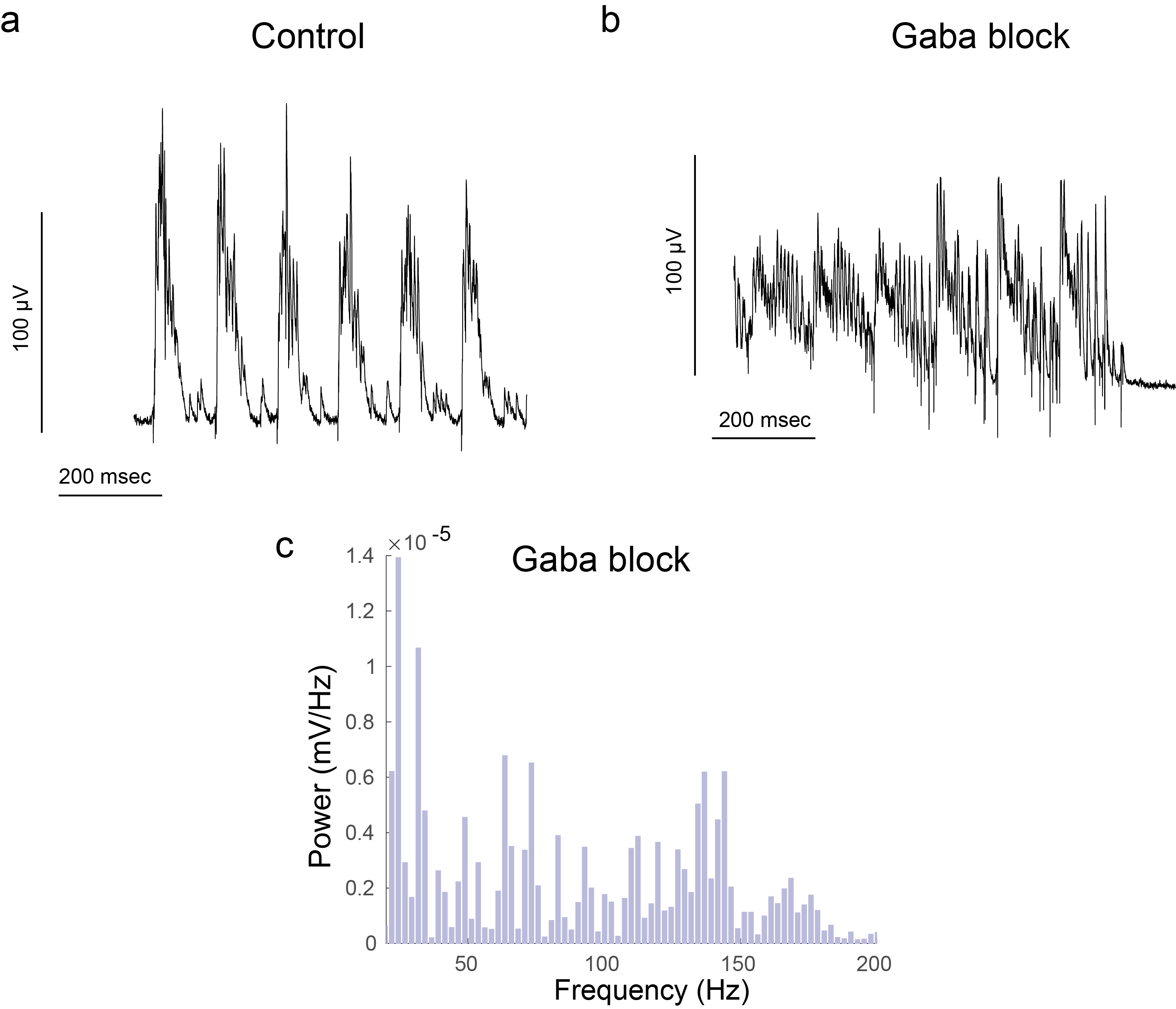

**Supplementary Figure 4 (related to Figure 3) simulation of the typical “Gabazine” experimental protocol** **(a)** Representative simulated LFP trace from the structured CA1 network under baseline control conditions, demonstrating the spontaneous emergence of discrete, organized SWR events. **(b)** Simulated LFP trace following the computational equivalent of pharmacological GABA blockade, achieved by setting all GABA synaptic conductances to zero. The network fails to generate organized SWRs. **(c)** Power spectral density (PSD) estimate of the LFP trace under the GABA-block condition. The characteristic SWR spectral signature is completely abolished. These simulations confirm that inhibitory synaptic conductances are required for the emergence of the SWR rhythm, preventing the network from collapsing into non-functional, hyperexcitable activity.

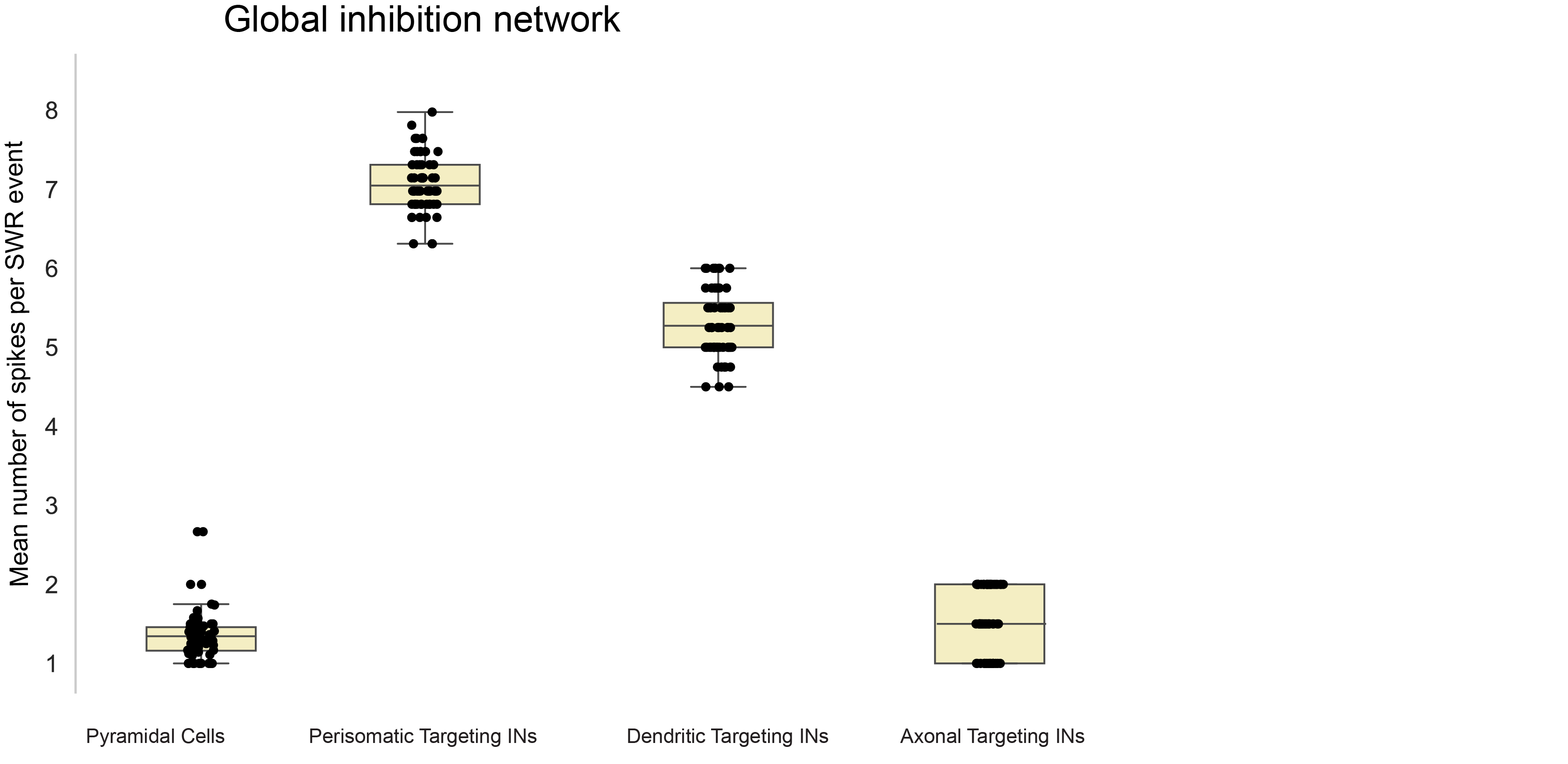

**Supplementary Figure 5 (related to Figure 4) Global inhibitory topology preserves neuronal participation and firing dynamics.** To ensure that the observed network-level effects are attributable to the redistribution of inhibitory connectivity rather than alterations in individual cellular excitability, we quantified the firing rates of each neuronal population under the global inhibitory topology configuration. The mean number of spikes per SWR event for all populations remained similar to those observed in the structured network model. These results confirm that the global inhibitory topology network maintains the characteristic physiological firing phenotypes required for SWR generation.

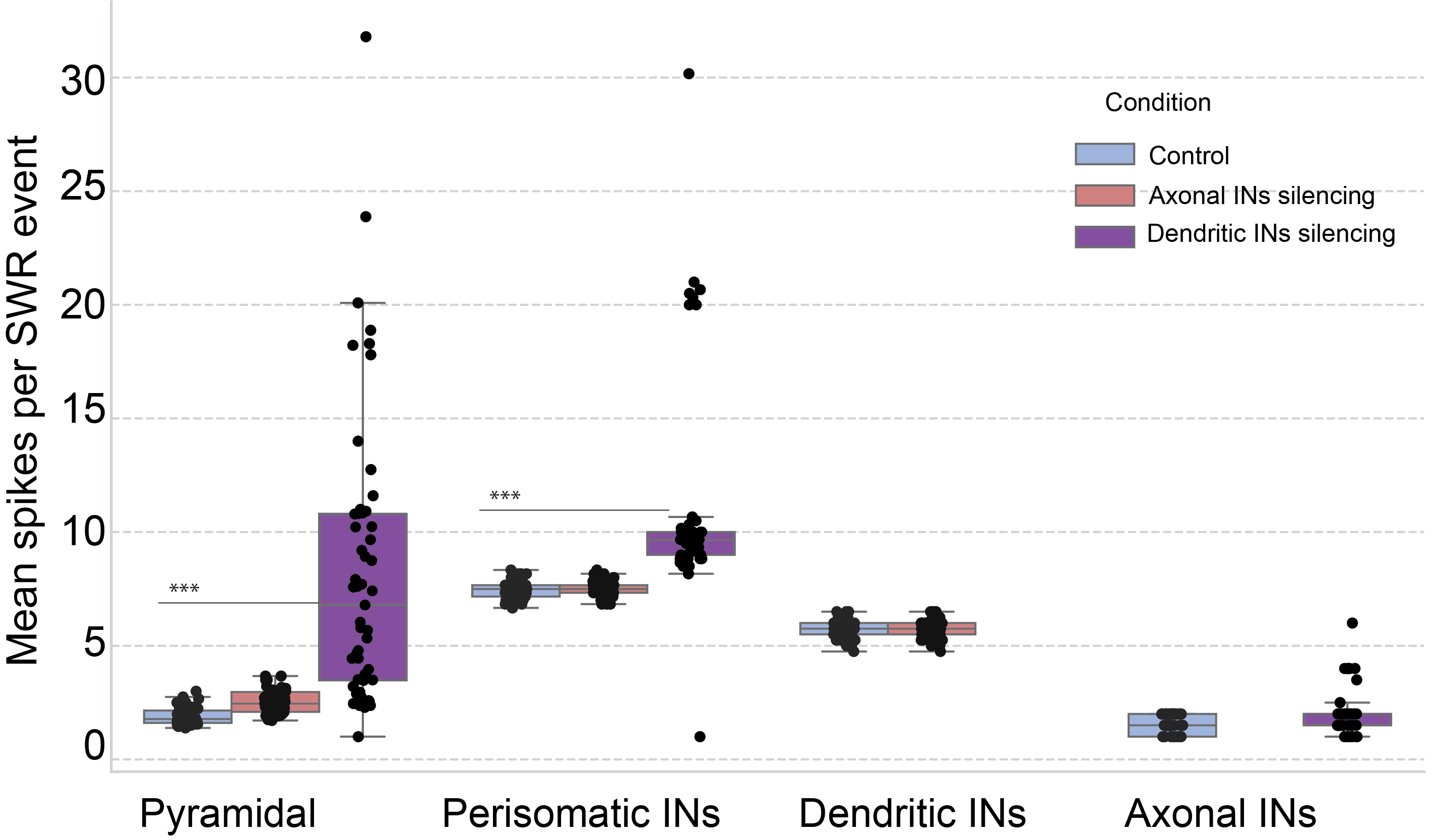

**Supplementary Figure 6 (related to Figure 6) Impact of targeted interneuron silencing on population-wide spiking activity.** Box plots illustrating the mean number of spikes per SWR event for each neuronal population under control (blue), axonal-targeting interneuron silencing (light red), and dendritic-targeting interneuron silencing (purple) conditions. The data demonstrate that dendritic-targeting interneuron silencing triggers a profound disinhibitory surge, manifested as a significant increase in the firing activity of both pyramidal cells and perisomatic-targeting interneurons. Conversely, silencing of axonal-targeting interneurons induces only modest changes in pyramidal cell spiking, with no significant alterations in the firing patterns of other inhibitory populations.

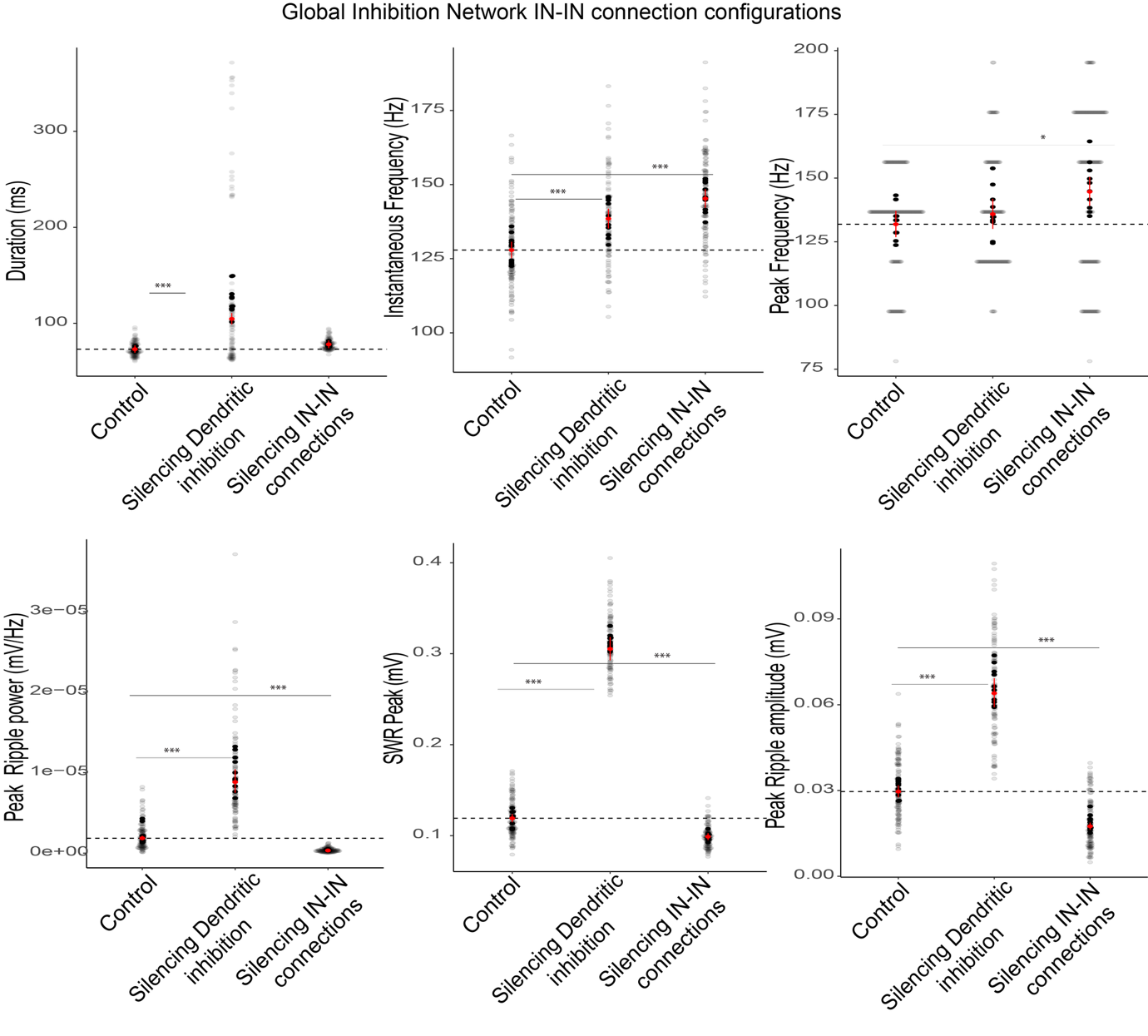

**Supplementary Figure 7 (related to Figures 6 & 7) Regulatory roles of inhibitory subpopulations and IN-IN motifs are conserved under global inhibitory topology.** To confirm that the functional roles of specific interneuron classes and interneuron-to-interneuron (IN-IN) connectivity are not dependent on the local spatial organization of the inhibitory network, we replicated the silencing protocols from Figures 6 and 7 within the global inhibitory topology configuration. Panels quantify the impact of silencing dendritic-targeting interneurons and total IN-IN connectivity on SWR duration, instantaneous frequency, peak frequency, peak ripple power, SWR peak amplitude, and peak ripple amplitude. Data represent n = 120 analyzed SWR events per experimental condition (pooled across both recording electrodes), extracted across 10 independent, randomized simulation trials. Statistical significance was evaluated using generalized linear mixed-effects models (***p < 0.001). These results demonstrate that dendritic-targeting interneurons and reciprocal IN-IN interactions exert the same spectral and magnitude-based control over SWR dynamics regardless of inhibitory topology.

**Supplementary Tables**

**Supplementary Table 1. Connectivity architecture of the CA1 Network.**

| **Presynaptic population** | **Postsynaptic population** | **Number of presynaptic partners per postsynaptic neuron** |
| --- | --- | --- |
| CA3 Schaffer collateral | Pyramidal cells | 270 |
| CA3 Schaffer collateral | Perisomatic-targeting interneurons | 200 |
| CA3 Schaffer collateral | Dendritic-targeting interneurons | 200 |
| CA3 Schaffer collateral | Axonal-targeting interneurons | 200 |
| Pyramidal cells | Pyramidal cells | 2 |
| Pyramidal cells | Perisomatic-targeting interneurons | 15 |
| Pyramidal cells | Dendritic-targeting interneurons | 15 |
| Pyramidal cells | Axonal-targeting interneurons | 13 |
| Perisomatic-targeting interneurons | Pyramidal cells | 17 |
| Dendritic-targeting interneurons | Pyramidal cells | 17 |
| Axonal-targeting interneurons | Pyramidal cells | 14 |
| Perisomatic-targeting interneurons | Perisomatic-targeting interneurons | 2 |
| Perisomatic-targeting interneurons | Dendritic-targeting interneurons | 4 |
| Perisomatic-targeting interneurons | Axonal-targeting interneurons | 2 |
| Dendritic-targeting interneurons | Perisomatic-targeting interneurons | 3 |
| Dendritic-targeting interneurons | Dendritic-targeting interneurons | 3 |
| Dendritic-targeting interneurons | Axonal-targeting interneurons | 3 |

Based on experimental reports from ^1–4^

**Supplementary Table 2. Membrane Properties**

| **Parameter** | **Pyramidal cells** | **Perisomatic-targeting interneurons** | **Axonal-targeting interneurons** | **Dendritic-targeting interneurons** |
| --- | --- | --- | --- | --- |
| **Passive membrane properties** |  |  |  |  |
| Membrane capacitance, (µF/cm²) | 1.0 | 1.4 | 1.4 | 1.4 |
| Axial resistance, (Ω·cm) | 150 | 100 | 100 | 100 |
| Leak conductance (S/cm²) | 3.33× 10⁻⁵ | 6.0 × 10⁻⁵ | 1.3 × 10⁻⁴ | 1.02 × 10⁻⁴ |
| **Sodium currents** |  |  |  |  |
| Fast Na⁺ conductance (S/cm²) | 1.12×10⁻² soma  1.26×10⁻² proximal apical dendrites  1.26×10⁻² medial apical dendrites  7.0×10⁻³ distal apical dendrites  7.0×10⁻³ basal dendrites 2.0×10⁻¹ axons | 0.169 | 0.135 | 0.090 |
| **Potassium currents** |  |  |  |  |
| Fast delayed rectifier K⁺ conductance (S/cm²) | 7.0×10⁻⁴ soma  8.68×10⁻⁴ proximal dendrites  8.68×10⁻⁴ medial apical dendrites  8.68×10⁻⁴ distal apical dendrites  8.68×10⁻⁴ basal dendrites  2.0×10⁻² axons | 0.011 | 0.0195 | 0.018 |
| A-type K⁺ (proximal) | 7.5×10⁻³ soma  1.5×10⁻² proximal dendrites | 0.00015 | 0.001 | 0.002 |
| A-type K⁺ (distal) | 3.0×10⁻² proximal and medial apical dendrites  4.5×10⁻² distal apical dendrites | -- | -- | -- |
| M-current | 6.0×10⁻² soma  6.0×10⁻² proximal apical dendrites  6.0×10⁻² medial apical dendrites  6.0×10⁻² distal apical dendrites  6.0×10⁻² proximal dendtites | — | — | — |
| Slow Ca²⁺-activated K⁺ current (sAHP) | 1.5×10⁻³ soma  5×10⁻⁴ dendrites | — | — | — |
| Medium Ca²⁺-activated K⁺ current (mAHP) | 4.54×10⁻¹ soma  3.3×10⁻² apical  4.1×10⁻³ basal | 0.002 | 0.002 | 0.002 |
| **Calcium currents** |  |  |  |  |
| L-type Ca²⁺ current | 7×10⁻⁴ soma  3.16×10⁻⁵ dendrites | 0.005 | 0.005 | 0.005 |
| T-type Ca²⁺ current | 5×10⁻⁵ soma  1×10⁻⁴ dendrites | — | — | — |
| R-type Ca²⁺ current | 3×10⁻⁴ soma  3×10⁻⁵ basal and apical dendrites | — | — | — |
| **Other currents** |  |  |  |  |
| Hyperpolarization-activated current | 5×10⁻⁵ soma  1×10⁻⁴ proximal apical  2×10⁻⁴ medial apical  3.5×10⁻⁴ distal apical  5×10⁻⁵–1×10⁻⁴ basal | — | — | — |
| Intracellular Ca²⁺ buffering | Present | — | — | — |

Based on simulation from ^1–3^

**Supplementary Table 3. Synaptic properties**

| **Presynaptic population** | **Postsynaptic population** | **Receptor** | **Synaptic target** | **Synaptic weight (g_{max})** |
| --- | --- | --- | --- | --- |
| **External CA3 Schaffer collateral input** |  |  |  |  |
| CA3 | Pyramidal cells | AMPA/NMDA | Apical and basal dendrites | 3.0 × 10⁻⁴ |
| CA3 | Perisomatic-targeting interneurons | AMPA | Dendrites | 2.25 × 10⁻⁴ |
| CA3 | Dendritic-targeting interneurons | AMPA | Dendrites | 5.70 × 10⁻⁴ |
| CA3 | Axonal-targeting interneurons | AMPA | Dendrites | 1.87 × 10⁻⁴ |
| **Internal excitatory CA1 connectivity** |  |  |  |  |
| Pyramidal cells | Pyramidal cells | AMPA/NMDA | Dendrites | 4.20 × 10⁻³ |
| Pyramidal cells | Perisomatic-targeting interneurons | AMPA | Dendrites | 7.70 × 10⁻⁴ |
| Pyramidal cells | Dendritic-targeting interneurons | AMPA | Dendrites | 1.90 × 10⁻³ |
| Pyramidal cells | Axonal-targeting interneurons | AMPA | Dendrites | 4.00 × 10⁻⁵ |
| **Internal inhibitory CA1 connectivity** |  |  |  |  |
| Perisomatic-targeting interneurons | Pyramidal cells | GABA(_A) | Soma and proximal dendrites | 1.08 × 10⁻³ |
| Dendritic-targeting interneurons | Pyramidal cells | GABA(_A) | Apical and basal dendrites | 2.75 × 10⁻³ |
| Dendritic-targeting interneurons | Pyramidal cells | GABA(_B) | Apical and basal dendrites | 2.75 × 10⁻³ |
| Axonal-targeting interneurons | Pyramidal cells | GABA(_A) | Axon initial segment | 1.24 × 10⁻² |
| Perisomatic-targeting interneurons | Perisomatic-targeting interneurons | GABA(_A) | Dendrites | 1.76 × 10⁻³ |
| Perisomatic-targeting interneurons | Dendritic-targeting interneurons | GABA(_A) | Dendrites | 3.19 × 10⁻³ |
| Perisomatic-targeting interneurons | Axonal-targeting interneurons | GABA(_A) | Dendrites | 1.32 × 10⁻⁴ |
| Dendritic-targeting interneurons | Perisomatic-targeting interneurons | GABA(_A) | Dendrites | 9.90 × 10⁻³ |
| Dendritic-targeting interneurons | Dendritic-targeting interneurons | GABA(_A) | Dendrites | 5.61 × 10⁻⁴ |
| Dendritic-targeting interneurons | Axonal-targeting interneurons | GABA(_A) | Dendrites | 6.60 × 10⁻⁴ |

Based on simulation from ^1–3^

**Supplementary Table 4. Electrophysiological properties**

| **Electrophysiological property** | **Pyramidal cells** | **Perisomatic-targeting interneurons** | **Axonal-targeting interneurons** | **Dendritic-targeting interneurons** |
| --- | --- | --- | --- | --- |
| Rheobase (pA) | 200 | 130 | 280 | 201 |
| Input resistance (MΩ) | 90 | 190 | 78 | 100 |
| Resting membrane potential (mV) | -65 | -68 | -68 | -68 |

Calibrated based on^2,5,6^
